# Activation of developmental transcription factors using RNA technology promotes heart repair

**DOI:** 10.64898/2026.01.16.700013

**Authors:** Riley J. Leonard, Mason Sweat, Steven Eliason, William Kutschke, Brad A. Amendt

## Abstract

Ischemic injury and adverse post-infarction myocardial remodeling are major causes of heart failure. We previously reported that microRNA (*miR)-200c* inhibition in murine embryos increased cardiogenic transcription factors (TFs) *Tbx5*, *Gata4*, and *Mef2c* to activate an immature cardiomyocyte cell state, suggesting *miR-200* inhibition as a therapy for cardiac repair. We performed permanent ligation of the left anterior descending artery (a severe myocardial infarct, MI) on *PMIS-miR-200c* (inhibition of *miR-200c; PMIS-C*), *PMIS-A* (inhibition of *miR-200a*) and wildtype adult mice. Echocardiographic left ventricular (LV) ejection fraction (EF) at 3 WPI (weeks post-injury) was 22% ± 4.31% (WT) but increased to 56% ± 4.25% (*PMIS-C*) (p ≤ 0.0001). Post-infarction LV chamber dilation was reversed in *PMIS-C* mice compared to WT, and trichrome staining showed a decrease in fibrosis 3 WPI. By 9 WPI, *PMIS-C* heart function was like that of WT mice before injury. Tbx5, Gata4, Mef2c, and Isl-1 were increased after MI in *PMIS-C* hearts. *PMIS-C* mice recover cardiac function and reverse ischemic pathology of acute cardiac injury in adult mice. Inhibition of *miR-200c* activates several important pathways in heart development and repair mechanisms after an MI in adult hearts. The *PMIS-miR-200c* transgenic mice demonstrate an important role for *miR-200c* in regulating heart repair after ischemic injury.

**Novelty and Significance:** *What is known?:* **\***The *microRNA-200* (*miR-200*) family targets several heart factors *in vitro*. \**miR-200c* inhibition was shown to protect cardiomyocytes in a myocardial ischemia-reperfusion injury, myocardial cellular model. \**miR-200* may play a role in cardiovascular fibrosis, however there are no *in vivo* reports of the role *miR-200* plays in heart repair.

*What New Information Does This Article Contribute?:* \**PMIS-miR-200c* transgenic mice reveal a role for *miR-200c* inhibition in rapid repair of the heart after a myocardial infarct (MI). After an MI, *miR-200c* expression increases, to levels observed during early heart development. **\***Inhibition of *miR-200c* allows for expression of Tbx5, Gata4, Pitx2, Mef2c, Yap, Nppa and Sox5 factors to repair the heart after ischemic injury. \**PMIS-miR-200c* mice have increased cardiomyocyte proliferation and reduced cardiac fibroblasts resulting in decreased fibrosis. *Heart function in *PMIS-miR-200c* mice is significantly restored 3-weeks post-MI. While microRNAs have been extensively studied in heart development and ischemic injury, little is known about the *miR-200* family in the cardiovascular system. In other cell types and systems, *miR-200* is upregulated under oxidative stress and hypoxia. *miR-200c* targets Zeb1, eNOS, Sirt1 and Fox01 to regulate cell growth and arrest, apoptosis and senescence in other tissues. *miR-200* members are increased in response to ischemia, but this has not been evaluated in the heart. We show a direct effect of *miR-200c* inhibition and decreased fibrosis in the MI heart. The *miR-200* family targets stem cell factors such as Sox2, Klf4, and Bmi1 and our recent sn-RNA multiomics analyses of *PMIS-miR-200c* mice revealed de-differentiated or immature cardiomyocytes. Thus, inhibition of *miR-200c* reactivates transcription factors after an MI, important for cardiomyocyte renewal. This research demonstrates how inhibition of *miR-200* regulates cardiac function after an MI.

## INTRODUCTION

Cardiovascular disease (CVD) is one of the leading causes of morbidity and mortality in the United States, and around the world ^1^. CVDs encompass a range of pathological changes to the heart, ultimately leading to a deterioration of function, leading to heart failure. Myocardial infarct/ischemic injury is a common CVD, in which occlusion of a coronary artery prevents perfusion of the working myocardium, resulting in cardiomyocyte (CM) death ^1^. As a result, the heart undergoes dilation and a fibrotic scar is formed in the space once occupied by the CMs, leading to a decrease in heart function ^2^. Progression of the pathology of ischemic injury can lead to heart failure and death (reviewed in ^2^).

The adult mammalian heart cannot induce proliferation, resulting in the loss of working myocardium following an injury (reviewed in ^3,4^. However, in lower vertebrates, adult cardiac regeneration can occur ^5^. Additionally, the neonatal mammalian heart can repair, but the ability diminishes quickly following birth ^6,7^. The mechanism of adult cardiac regeneration can be achieved through expression of positive cell cycle regulators ^8,9^, cardiomyocyte de-differentiation ^10–12^, cardiac fibroblast programming ^11,13–15^, and cell-based therapies ^16–18^. Molecular mechanisms of successful approaches have targeted Cyclin D2 ^9^, cardiogenic transcription factors ^12^, Hippo pathway regulators ^7^, and microRNAs (miRs) ^8,10,13,19–26^. A recent commentary discussed the potential role of the *miR-200* family in cardiovascular diseases, noting a lack of research on heart function ^27^. The *miR-200* family has been implicated in SMC (smooth muscle cell) dedifferentiation during injury-induced vascular remodeling ^28^. Genome analysis has predicted that *miR-200c* expression, which targets ZEB1/2, plays a role in the pathology of patients with bicuspid aortic valves ^29^. In a myocardial ischemia-reperfusion injury model *miR-200c* inhibition *reduced* cardiomyocyte death and apoptosis Zhang, 2022]. Our research has identified several stem cell factors, transcription factors and developmental processes regulated by the *miR-200* family in mice ^30–34^. We have previously identified factors regulating heart development in the *PMIS-miR-200* (<u>P</u>lasmid-based <u>m</u>icroRNA Inhibitor <u>S</u>ystem, targeting *miR-200*) transgenic mice ^30,34^.

We hypothesized that inhibition of *miR-200c* activity would result in cardiomyocyte dedifferentiation sufficient to ameliorate post-infarction decompensation after ischemic injury in adult mice. In this report, we demonstrate that inhibition of *miR-200c* (*PMIS-miR-200c*, *PMIS-C* transgenic mice) can promote rapid and sustained adult cardiac repair following permanent ligation of the left anterior descending (LAD) artery (MI). We found that *miR-200c* was upregulated following ischemic injury and that transgenic mice with inhibited *miR-200c* expression (*PMIS-C* mice) had a significant increase in left ventricle ejection fraction (LVEF) and decreased LV dilation by 3 weeks post-injury (3 WPI), which continued to improve by 9WPI. Mechanistically, inhibiting *miR-200c* increases the expression of cardiogenic transcription factors (TFs) *Tbx5*, *Gata4*, and *Mef2c* and inhibition of the Hippo pathway, with an increase in activated Yap1 in *PMIS-C* hearts. We observed Nppa+/Sox5+ border zone CMs at 1 WPI, which we identified as markers of an immature CM cell state, unique to the *PMIS-C* heart ^34^. In the *PMIS-C* heart, CM proliferation was significantly increased, while non-cardiomyocyte proliferation was decreased compared to WT. Overall, this study provides evidence that inhibiting *miR-200c* can activate an immature state of CMs, leading to rapid heart repair following a severe MI in adult mice.

## METHODS

### Animals

Mice were housed and handled in accordance with guidelines established by the University of Iowa Institutional Animal Care and Use Committee (IACUC). All experimental techniques were approved according to University of Iowa IACUC guidelines. Male and female mice were used in the study. The construction of *PMIS* inhibitor mice were previously described ^30^.

### Sex as a biological variable

Our study examined male and female animals, and similar findings are reported for both sexes.

### Murine myocardial infarct model

The surgeon performing the LAD procedure was blinded as to the genotype of the mice in all experiments. Murine myocardial infarct surgical procedure was performed as previously described ^35,36^. Briefly, male and female mice at 8-10 weeks of age were anesthetized with isoflurane (5% inhaled). The animals were intubated and ventilated with 2% isoflurane/98% O_2_ using a Harvard Apparatus Model 687 mouse ventilator. A left thoracotomy was performed, the heart was exposed, and the pericardium was removed. A suture was passed underneath the left anterior descending (LAD) branch of the coronary artery ∼3 mm from the tip of the left atrium along the anterolateral border of the heart (approximately mid-LAD coronary artery), and a surgical knot was tied to occlude the coronary artery. Successful ligation of the artery was confirmed by observing a blanching of the myocardium. At the end of the procedure, a chest tube (28-gauge, venal catheter) was placed between the fourth and fifth ribs, and then the chest wall was closed. The control group had a similar procedure, with the passage of a suture under the LAD artery but without occlusion. The infarct size of MI mice was measured based on echocardiography.

### Echocardiography

The technicians performing the echocardiograms were blinded to the genotype of all mice during all recordings. Transthoracic echocardiograms in conscious minimally sedated mice were performed in the University of Iowa Cardiology Animal Phenotyping Core Laboratory using a Vevo 2100 Imager (VisualSonics, Toronto, ON, Canada), as previously described ^35,36^. The anterior chest was shaved, and the prewarmed ultrasonic gel was applied. Two-dimensional images were acquired in LV short- and long-axis planes with a 40-MHz linear array probe, yielding 200 frames/s. LV mass, volume, and ejection fraction (EF) were calculated with the biplane area-length method, as previously described ^37^. Regions demonstrating akinesis or dyskinesis were visually identified, traced, and expressed as percentages of total LV end-diastolic silhouette.

### RNA isolation from tissue

RNA was isolated from freshly dissected tissues or cells using Trizol reagent (Thermo Fisher). RNA quality was assessed by gel electrophoresis prior to reverse transcription (RT) applications. 0.5-1μg of total RNA was used in an RT reaction with the 5x primescript RT kit (Takara Bio). cDNA was diluted 1:5, and 1μl was used per qPCR reaction. qPCR reactions were normalized to *β-actin* levels. For miRs, RT was performed using the Qiagen miRScript kit. The resultant cDNA was diluted 1:5 and 1μl was used per qPCR reaction. qPCR probes for specific miRs were generated, as well as the normalization set for murine U6. PCR products were examined by melting curve analysis and the specificity of the qPCR products were confirmed by sequencing. Fold changes were calculated using the 2^−ΔΔCT^ method. Primers are listed in table 1.

**TABLE 1.**
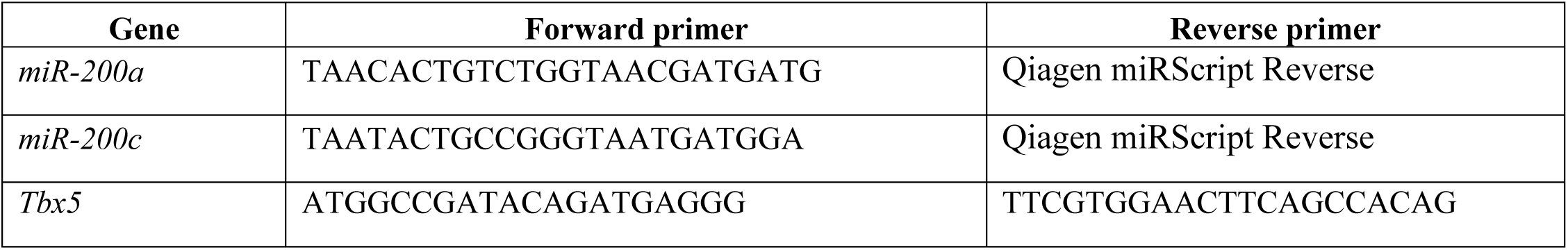
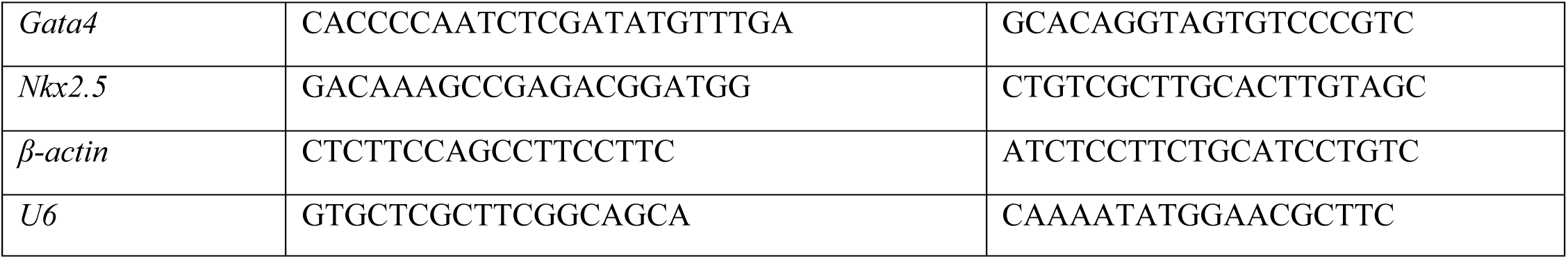
Primers for qPCR analyses.

### Paraffin embedding, sectioning, and staining

After embryo isolation and dissection, embryos were fixed for 30 min at 4°C in 4% paraformaldehyde (PFA) and dehydrated through an ethanol gradient. Dehydrated tissues were pre-cleared with xylene and incubated with three changes of paraffin, and then embedded into blocks. Paraffin-embedded tissues were sectioned and mounted on slides. Prior to staining, slides were de-paraffinized and rehydrated through a reverse ethanol gradient. Hematoxylin and eosin (H&E) and Trichrome staining, which has been previously described, were performed to assess tissue morphology ^38^.

### Histology and fluorescent immunohistochemistry

Immunofluorescence (IF) was performed as previously described ^32^. In brief, sections were rehydrated in xylene and ethanol series and then incubated in Block Buffer (1% BSA, 2.5% goat serum, 1% phosphate-buffered saline with 0.25% Triton X-100, PBST) for 30 min at room temperature, washed, followed by overnight incubation at 4°C with primary antibody (Ab). After the incubation, the slides were treated with goat anti-mouse Alexa-488 or Alexa-594 and/or goat anti-rabbit Alexa-488, Alexa-555, Alexa-594 or Alexa-647 labeled secondary antibody (Invitrogen) at a concentration of 1:300 for 60 min. Each antibody incubation was followed by 3–6 PBST washes. Nuclei were counterstained with DAPI prior to adding coverslips with mounting media (Vector Laboratories). For subsequent IF, primary rabbit Ab was added for overnight incubation at 4°C. Sections were incubated with secondary Ab for 60 mins. Sections were incubated in Blocking Buffer plus 2.5% rabbit serum for 30 min., followed by overnight incubation at 4°C with primary Ab. Sections were incubated with secondary Ab for 60 minutes and washed. Nuclei were counterstained with DAPI mounting media. Antibodies use in the experiments are listed in table 2.

**TABLE 2.** Antibodies used in the experiments.

|  | **Dilution** | **Source** |
| --- | --- | --- |
| Tbx5 | 1:100 | Abcam |
| cTnT (CT3) | 1:50 | DSHB-Lin, J.J.-C. |
| cTnT | 1:500 | Proteintech |
| Isl1 | 1:100 | Abcam |
| Nppa | 1:500 | Proetintech |
| Sox5 (PCRP-SOX5-1E3) | 1:25 | DSHB-NIH |
| BrdU | 1:100 | Abcam |
| Vim | 1:100 | Abcam |
| PDGFRα | 1:100 | Proteintech |
| αSMA | 1:1000 | Abcam |
| CD90/Thy1 | 1:50 | DSHB-NIH |
| CC3 | 1:100 | Abcam |
| Collagen I | 1:100 | DSHB-DeBlas, Angel, L. |
| Fibronectin | 1:100 | DSHB-Wayner, E.A. |

### Western Blot

Immunoblots were performed as previously described ^32^. In brief, tissue or cell lysates were analyzed on SDS-PAGE gels. Adult cardiac tissue was lysed in RIPA buffer. Following electrophoresis, the protein was transferred to a PVDF membrane (Millipore), immunoblotted, and detected with an HRP-conjugated secondary antibody and ECL reagents (GE Healthcare/Amersham Biosciences). The following polyclonal antibodies were used to detect the proteins: Mef2c (1:1000) (Cell Signaling), Tbx5 (1:500) (Thermo Fisher), Gata4 (1:1000) (Santa Cruz), Yap1 (1:2000) (Abcam), pYap1 (1:2000) (Abcam) and Gapdh (1:8000) (Santa Cruz).

### Statistical methods

P values were calculated using two-tailed t-tests with a minimum of three biological replicates for each group. Error bars are presented as +/- standard error of the mean in every figure. *, indicates a P value of <0.05, ** <0.01, *** <0.001, ****<0.0001.

## RESULTS

### Permanent ligation of the left anterior descending artery (LAD) and myocardial infarct recovery in *PMIS-miR-200* mice

During murine embryonic cardiac development, *miR-200c* is expressed at high levels compared to adults (Fig. 1A) ^34^. As CMs reactivate an early developmental gene regulatory network (GRN) following injury ^10–12^, we hypothesized that expression of *miR-200c,* which is an inhibitor of reactivation of embryonic gene expression ^34^, would also increase. Indeed, expression of *miR-200c* significantly increased post-MI (Fig. 1A), suggesting the family has a role in the injured adult heart.

**Fig. 1.**
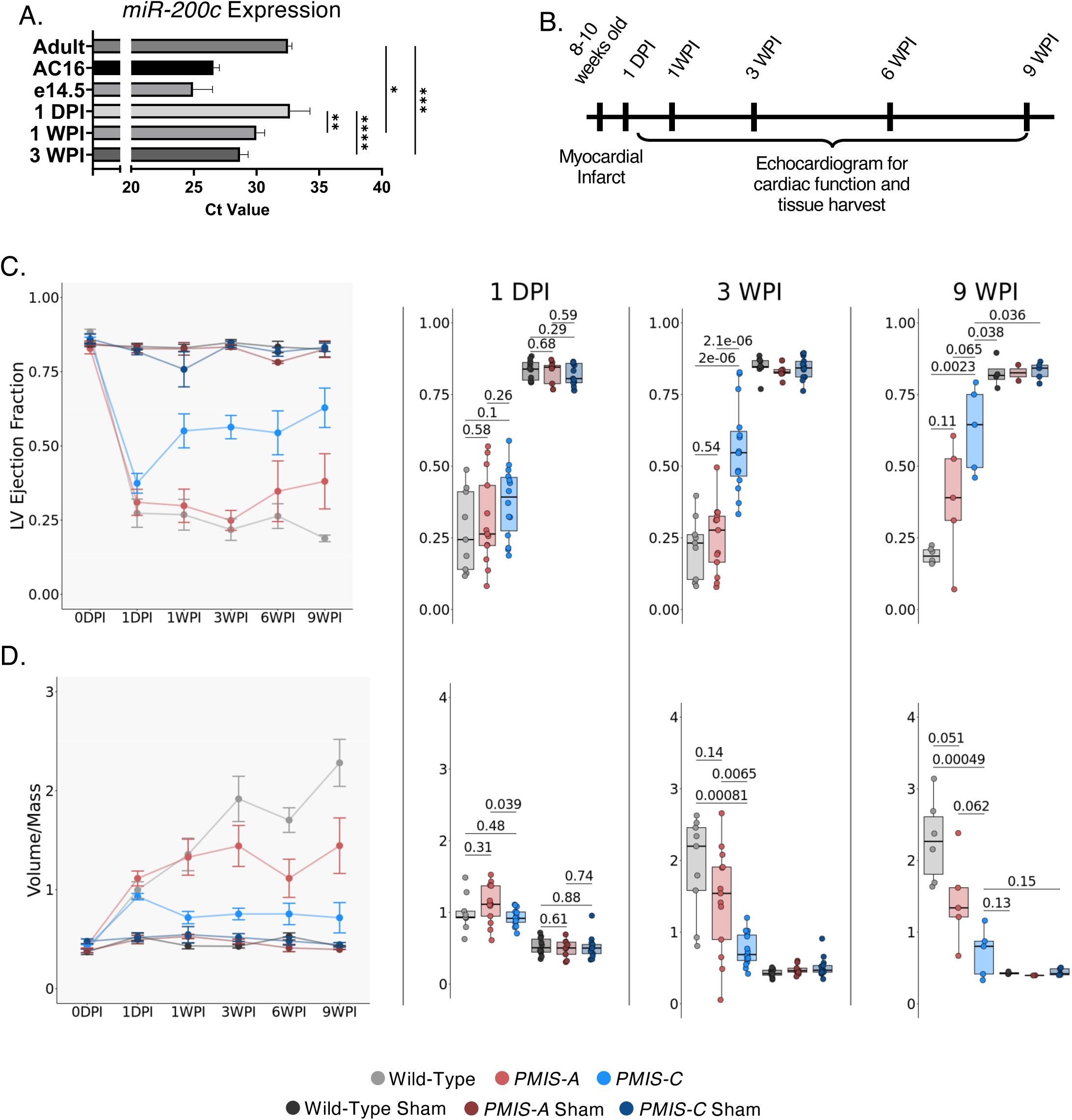
*PMIS-C* mice show functional recovery post-MI. **A)** Expression of *miR-200c* in WT adult, embryonic and LAD injured hearts at 1 DPI (day post-injury), 1 WPI (week post-injury) and 3 WPI. N=4-7. Note expression is shown as direct Ct values and not fold change. Increased Ct values denote low expression. **B)** Timeline of LAD surgery and follow-up echocardiograms and molecular analysis. **C)** LVEF of WT, *PMIS-A*, and *PMIS-C* mice subject to LAD or sham procedure. LVEF was measured by echocardiogram at 1 DPI to 9 WPI. **D)** LV volume/mass of WT, *PMIS-A*, and *PMIS-C* mice subject to LAD or sham procedure to 9 WPI. The statistical test performed was a two-tailed t-test.

To investigate the hypothesis that *PMIS-miR-200* (*PMIS-miR-200a*, inhibits *miR-200a* and *miR-141*) and (*PMIS-miR-200c*, inhibits *miR-200b*, *miR-200c*, *miR-429*) mice will promote and sustain rapid cardiac repair following a myocardial infarct/ischemic injury (MI), we subjected 8–10-week-old mice to a permanent LAD ligation. This procedure is a model to study MI in rodents and other animal models ^39^. Following MI, adult mice were evaluated by echocardiogram at 1-day post-injury (1DPI). Mice with a ≥30% left ventricle (LV) ischemic zone fraction (IZ) and ≤50% LV ejection fraction (EF) were used for further studies, producing groups with equivalent severity of initial ischemic injury (Fig. S1A, B) ^39^. Kaplan-Meier death curves were analyzed for Wild-Type (WT), *PMIS-miR-200a (PMIS-A*) and *PMIS-miR-200c* (*PMIS-C*) mice subject to an MI (Fig. S1C). The *PMIS-A* and *PMIS-C* mice show a significant increase in survival as a first indicator of heart repair.

Echocardiograms were used to evaluate cardiac function at 1-, 3-, 6-, and 9-weeks post-injury (WPI) (Fig. 1B-D). Wild-Type, *PMIS-A* and *PMIS-C* mice had an average LVEF ≤50%, with no significant differences between the groups following LAD at 1 DPI (Fig. 1C). However, at 3 WPI, WT LVEF was 22% ± 4.31%, with significant changes at 56% ± 4.25% in *PMIS-C* mice (Fig. 1C; Data file S1). Moreover, the post-infarction LV chamber dilation (Vol/Mass ratio) was decreased in *PMIS-C* mice (Fig. 1D; Data file S1). The calculated delta infarct zone (ΔIZ) found a significantly decreased ratio in *PMIS-C* mice compared to controls, suggesting a decrease in the overall IZ of the LV (Fig. S2A; Data file S1). Measurements of end-systolic volume (ESV) and end-diastolic volume (EDV) support this finding for *PMIS-C* MI mice (Fig. S2B; Data file S1). There were no significant changes in heart rates of all mice (Fig. S2C; Data file S1).

We followed *PMIS-C* injured mice to 9 WPI to evaluate the long-term efficacy of the increased cardiac function. Echocardiogram analysis showed *PMIS-C* mice continued a trend of increased LVEF at 6 WPI (54% ± 7%) and 9 WPI (63% ± 7%) (Fig. 1C; Data file S1). LV dilation progressed over time in WT mice but reversed over time in *PMIS-C* mice (Fig. 1D). Overall, the data demonstrate that that *PMIS-C* mice have improved cardiac function by 3 WPI and continue to progress long-term. Functional recovery in *PMIS-C* hearts, observed by echocardiogram, suggests that inhibiting *miR-200c* expression is beneficial to ischemic heart repair and diminishes LV remodeling following MI.

### Post-MI fibrosis and scar size are decreased in *PMIS-C* mice

To determine that the LAD procedure resulted in myocardial cell death, we analyzed the expression of the apoptotic marker CC3 at 3 DPI (Fig. S3A). The results showed similar expression of CC3 in the LV free wall across all genotypes, indicating a loss of working myocardium below the ligation. We next performed histological serial sectioning and trichrome staining to determine the extent of fibrotic scarring. At 1 WPI, the WT, *PMIS-A*, and *PMIS-C* hearts show fibrotic scar in ischemic zone (IZ) and in the ischemic border zones (BZ) (Fig. S3B) ^40–42^. Trichrome staining, used to visualize the fibrotic scar, showed myocardium remodeling was present at 1 WPI (Fig. S3B). The magnified regions of the infarct zone (green box) and border zone (black box) show fibrosis (Fig. S3C, D). These results provide evidence that the LAD procedure results in cardiomyocyte cell death at a level sufficient to induce fibrotic remodeling.

Though scar formation is present at 1WPI (Fig. S3), by 3 WPI, *PMIS-C* hearts have a significantly reduced size of the fibrotic scar throughout serial sections compared to WT and *PMIS-A* mice (Fig. 2A, B, C). This reduced scar phenotype is seen consistently across individual *PMIS-C* hearts, providing evidence of the efficacy of inhibition of *miR-200c* towards cardiac repair in adult hearts. We show multiple hearts to demonstrate the overall decrease in fibrosis in *PMIS-C* hearts, compared to PMIS-A and WT, which adds to our rigor and reproducibility studies. Moreover, at 9 WPI, the *PMIS-C* hearts replicate and preserve reduced fibrotic scar size, observed at 3WPI (Fig. S4).

**Fig. 2.**
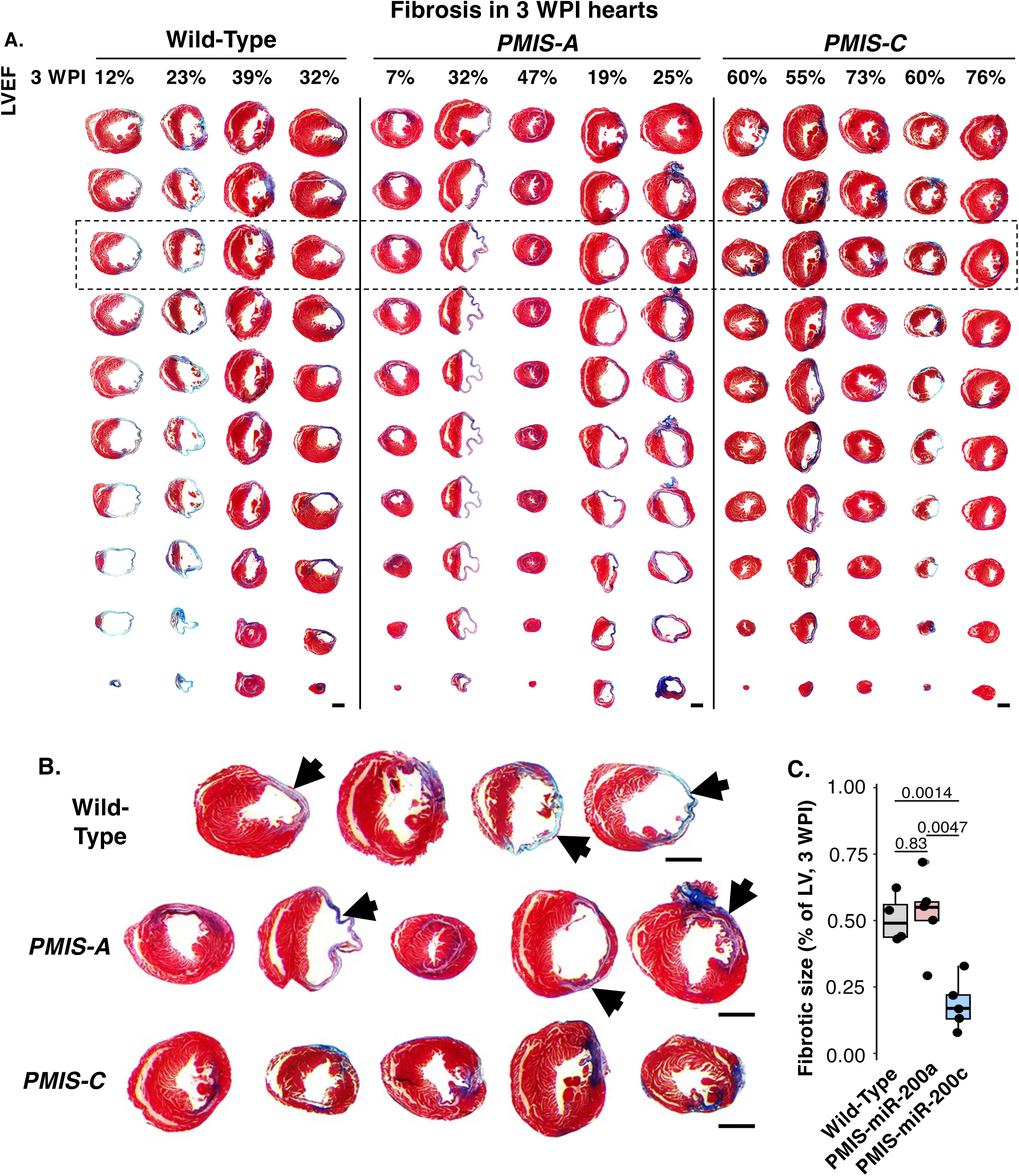
*PMIS-C* hearts have minimal scarring at 3 WPI. **A)** Trichrome staining of transverse ventricle sections of individual WT, *PMIS-A*, and *PMIS-C* subject to MI. Echocardiogram of LVEF for each heart is indicated at the top of the figure. **B)** Magnified images of individual hearts, arrows point to fibrosis (blue). Scale bar = 1mm. **C)** Quantitation of fibrotic size, % of LV, N=5.

Taken together, the echocardiogram and histological results indicate that inhibition of *miR-200c* promotes rapid and sustained activation of the myocardium after an MI.

### *PMIS-C* mice have an immature CM cell state marked by embryonic transcription factors that are present post-MI

During development, we identified cardiogenic transcription factors *Tbx5*, *Gata4*, and *Mef2c* as *miR-200c* mRNA targets during *in vivo* cardiogenesis ^34^. We used single-nuclei multiomics analysis to reveal an immature CM cell state present in the *PMIS-C* hearts ^34^. Based on those results, and previous studies showing the requirement for the TFs in regeneration ^12,43–45^, we hypothesized that these immature CMs in *PMIS-C* hearts could aid in adult cardiac regeneration. Indeed, at 1 and 3 WPI, we observed increased expression of cardiogenic TFs Tbx5, Gata4, and Mef2c in *PMIS-C* hearts compared to WT (Fig. 3A, B, C), consistent with our previously identified immature CM cell state ^34^. Additionally, the *PMIS-C* heart had a significant decrease in Hippo pathway activation, with a decrease in phosphorylated Yap protein and increased unphosphorylated Yap (Fig. 3A, B, C). These results support our previous data that inhibition of *miR-200c* increased specific transcription factors associated with an immature cell state, which, appears beneficial to cardiac repair (Fig. 3A,B,C) ^46,47^.

**Fig. 3.**
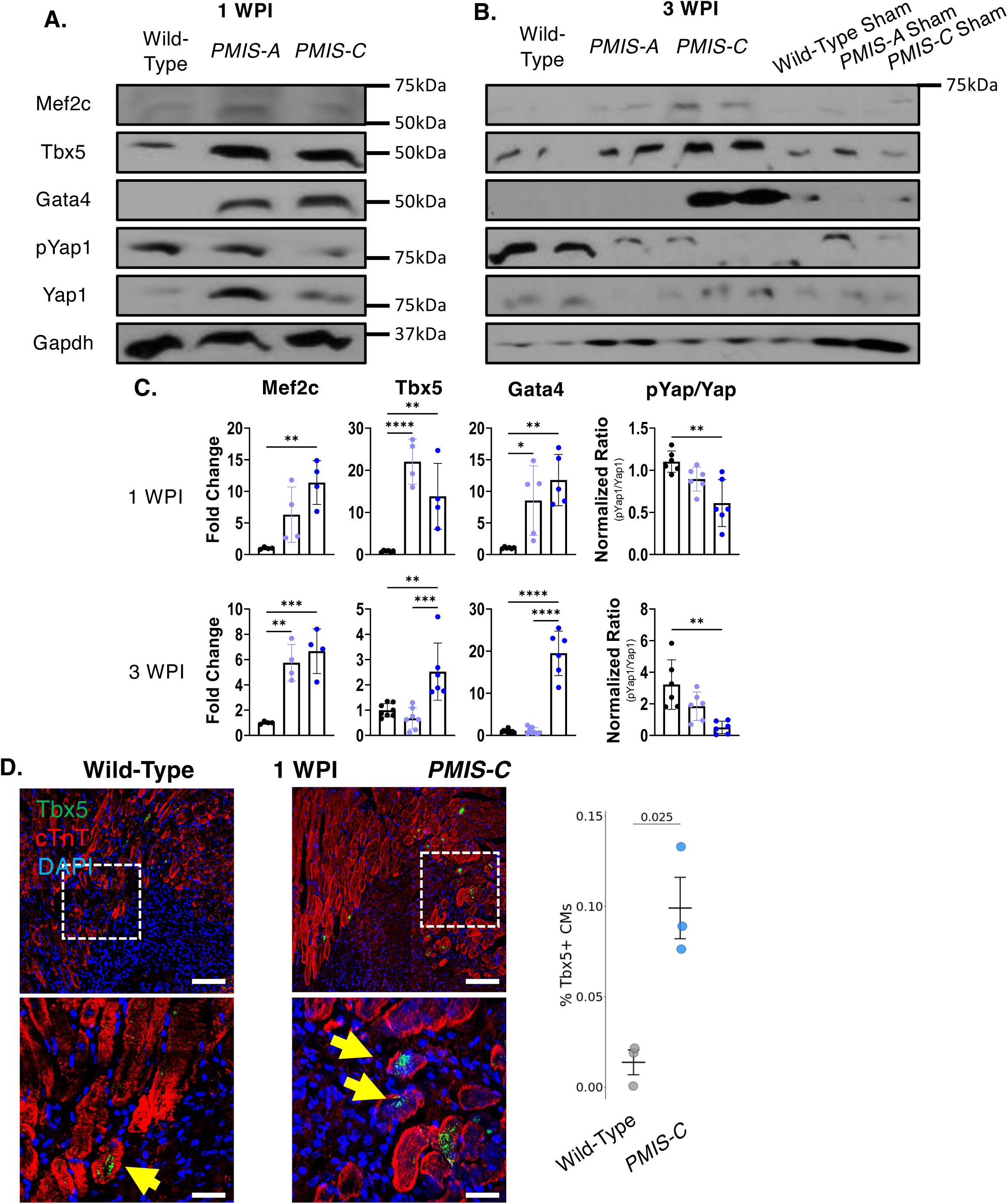
Cardiogenic TFs are increased in *PMIS-C* hearts following injury. **A-B)** Western blot for Mef2c, Tbx5, Gata4, pYap1, and Yap1 at 1 (A) and 3 WPI (B). Protein was isolated from the whole ventricular tissue. **C)** Quantification of Mef2c, Tbx5, and Gata4 normalized to Gapdh. The ratio of pYap/Yap at 1 WPI. The statistical test performed was a two-tailed t-test *=p≤0.05, **=p≤0.01, ***=p≤0.001, ****=p≤0.0001. **D)** IF stain of left ventricle sections in the BZ of WT and *PMIS-C* at 1 WPI for Tbx5. *PMIS-C* hearts have increased Tbx5+ CMs at the BZ (Yellow Arrows). Quantitation of Tbx5+ cells is shown for Wild-Type and *PMIS-C*. Scale bar = 50μm. The statistical test performed was a two-tailed t-test.

While the cardiogenic transcription factors were increased in *PMIS-C* hearts, we wanted to identify the expression in cardiomyocytes. At 1 WPI, Tbx5 showed expression in BZ cardiomyocytes, which the percentage of Tbx5+ CMs was significantly increased compared to Wild-Type (Fig. 3D). Furthermore, transcript analyses of whole hearts 1 WPI showed an increase in *Tbx5* and *Nkx2.5*, and 3 WPI showed increased *Tbx5*, *Gata4*, and *Nkx2.5* (Fig. S5).

Molecular analysis of the adult injured *PMIS-C* heart found increased expression of *miR-200c* targets and cardiogenic transcription factors Tbx5, Gata4, and Mef2c. These results suggest that, in both development and regeneration, *miR-200c* is targeting these factors to inhibit an immature CM state ^34^. Further, CMs with increased Tbx5 expression were observed at the BZ at 1 WPI, placing them in a location to facilitate adult cardiac repair. Following cardiac injury in adult mice, BZ cardiomyocytes are a population of de-differentiated, immature cells hypothesized to have a role in cardiac regeneration ^12,48–51^. Therefore, following ischemic injury, the activation of the immature CM cell state in *PMIS-C* hearts may lead to the reduced fibrosis and repair of the adult heart.

### The immature CM population is found at the border zone of *PMIS-C* adult ischemic hearts

To further validate the immature status of border zone (BZ) CMs, we assayed for Islet-1 (Isl1) (Fig. 4A), a known marker for cardiac progenitor cells ^4,52–55^. At the BZ, *PMIS-C* hearts had a significantly increased number of Isl1+ CMs (dashed boxed region, magnified in solid box) compared to WT (Fig. 4A). Next, we stained for Nppa and Sox5, which were identified as markers of the *PMIS-C* immature CMs ^34^. As expected, WT and *PMIS-C* BZ CMs showed Nppa expression in CMs (Fig. 4B) ^4^, however, CMs in the *PMIS-C* heart also expressed Sox5 at significantly higher levels (Fig. 4C). Additionally, *PMIS-C* CMs had a significant co-expression of Nppa and Sox5 (Fig. 4D, dashed boxed region magnified in the image below), suggesting the presence of *PMIS-C* immature cells at the border zone at 1 WPI. At 3WPI WT and *PMIS-C* BZ CMs showed Nppa and Sox5 expression in the CMs, but not in sham hearts (Fig. S6). Additionally, at 1 WPI the Remote Zone (RZ) of *PMIS-C*, showed a decrease in expression of Nppa and Sox5 (Fig. S7), suggesting the immature CM state is transient during adult cardiac repair. Overall, these results show that *PMIS-C* hearts, after ischemic injury, have an active and immature CM cell state that is reminiscent of *PMIS-C* embryonic cardiomyocytes ^34^.

**Fig. 4.**
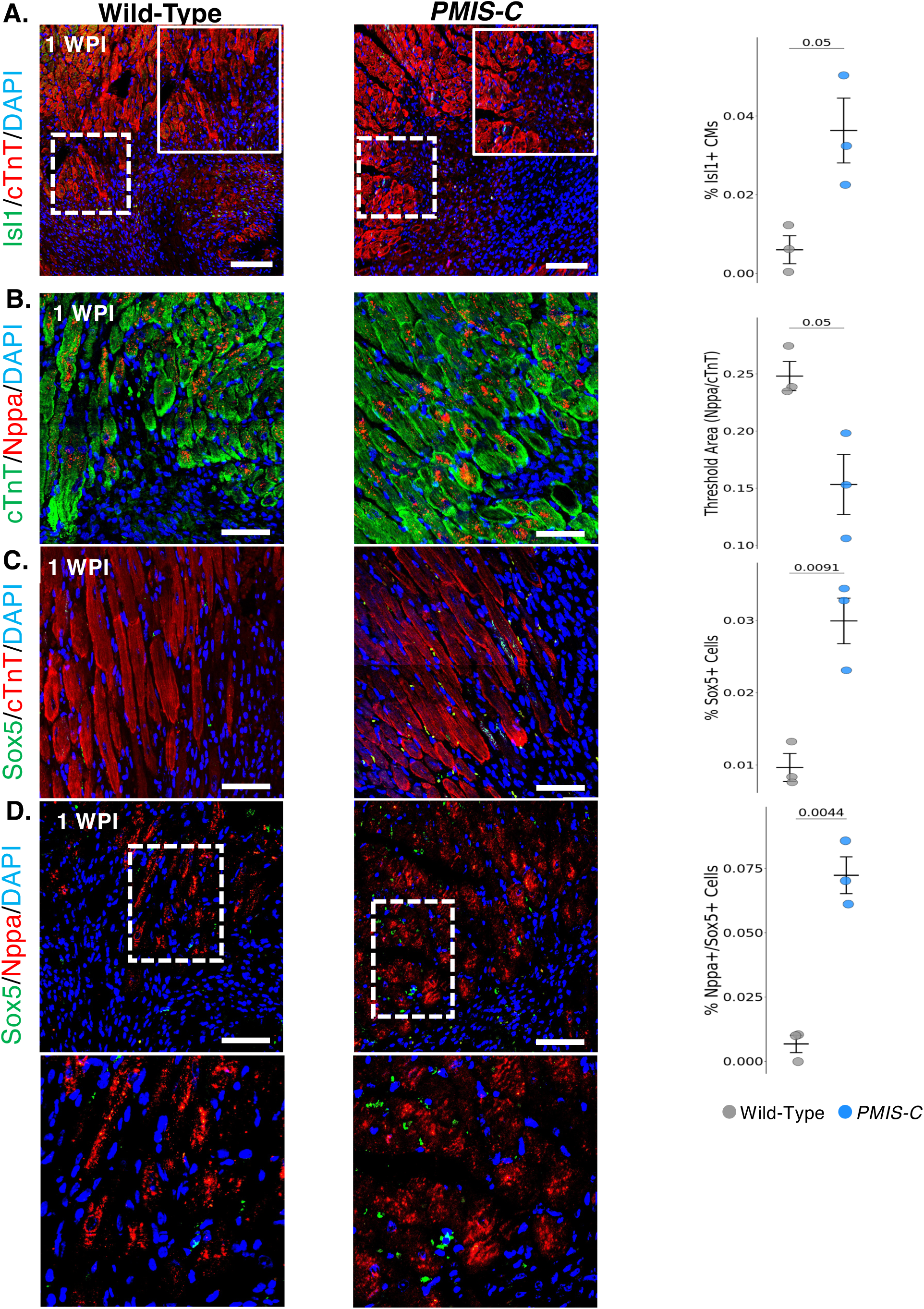
Cardiac immature markers are induced in *PMIS-C* CMs. **A)** IF stain of left ventricle sections in the BZ of WT and *PMIS-C* at 1 WPI (A) for Islet-1 (Isl1, green). *PMIS-C* hearts have increased Islet-1+ CMs at the BZ at 1WPI. **B-D)** IF stain of BZ CMs for Nppa (B, red), Sox5 (C, green), or both (D) in WT and *PMIS-C* injured hearts. Nppa is expressed in hearts subject to MI. Quantitation of Isl1+, Nppa/cTnT+, Sox5+ and Nppa/Sox5+ cells is shown. Scale bar = 50μm. The statistical test performed was a two-tailed t-test.

### In ischemic hearts, inhibition of *miR-200c* affects cell proliferation in a cell-type-specific manner

Our data showed that cell death and fibrotic scar formation occur in all genotypes post-injury, thus the injured heart requires new CMs to repair lost myocardium. To test the hypothesis that the new myocardium arises through CM proliferation, we performed a pulse-chase experiment by injecting thymidine analogs EdU and BrdU following MI (Fig. 5A). We analyzed hearts at 3 WPI, as *PMIS-C* mice show significantly increased function and reduced fibrosis at this time point. Analyses of EdU and BrdU incorporation in ischemic hearts found single positive (EdU+ or BrdU+) and dual positive (Edu+ and BrdU+) CMs in WT and *PMIS-C* hearts (Fig. 5B). Border zone CMs had a significant increase in BrdU+ labeled cells in *PMIS-C* versus WT, suggesting continuing CM proliferation in *PMIS-C* mice (Fig. 5B). The number of EdU+ CMs was unchanged, suggesting that EdU retaining CMs had proliferated and lost the label in both WT and *PMIS-C* mice, as expected. As these analogs incorporate into all proliferating cells, we also analyzed incorporation in cells within the infarct zone (Fig. 5C). Interestingly, in *PMIS-C* hearts, non-CMs (cTnT-) of the infarct zone had a significantly decreased number of EdU+ and BrdU+ cells compared to WT, suggesting inhibition of *miR-200c* reduces non-CM cell proliferation. The reduction in non-CM cell proliferation could be a factor in the observed cardiac repair in the *PMIS-C* adult heart.

**Fig. 5.**
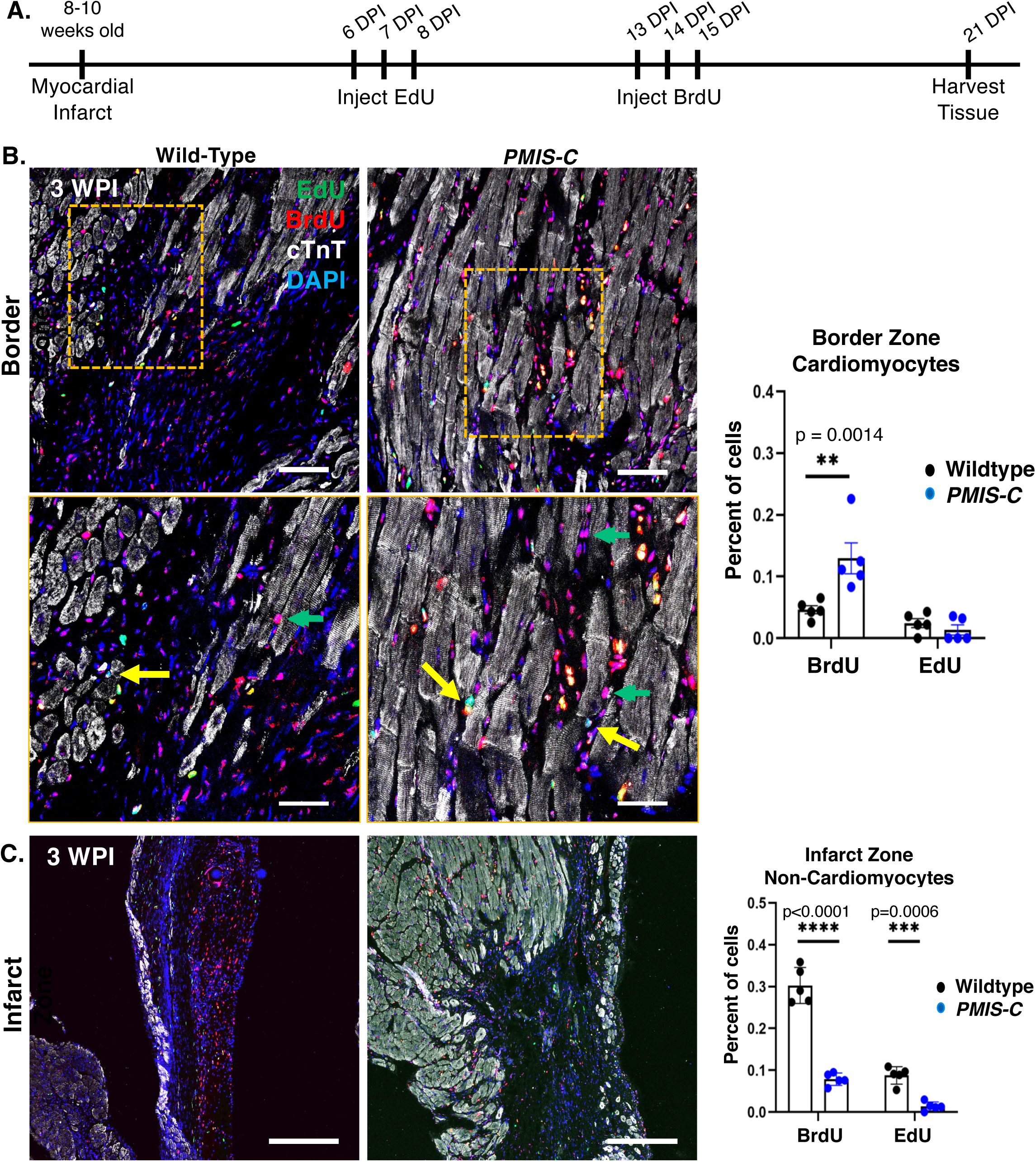
The proliferation of CMs may contribute to cardiac regeneration in *PMIS-C* hearts. **A)** Timeline of the pulse-chase experimental design to test proliferation rate/time following MI. **B)** IF stain of 3 WPI left ventricle sections for EdU (Green), BrdU (Red), and cTnT (White). Lower panel: The magnified image of the orange-boxed region above; images of the BZ show EdU+ (yellow arrow) and BrdU+ (green arrow) present in both hearts. There are more cTnT CMs in the *PMIS-C* border zone compared to WT, see quantitation. Scale bar = 100μm Bottom: Scale bar = 50μm. **C)** IF stain of 3 WPI left ventricle sections for EdU (Green), BrdU (Red), and cTnT (White). Representative images of the infarct zone show BrdU+ in non-CMs of the WT IZ. Non-CMs in *PMIS-C* hearts display a minimal number that incorporates either EdU or BrdU, see quantitation. Scale bar = 100μm. The statistical test performed was a two-tailed t-test.

### Fibroblast cell types may be differentially activated in *PMIS-C* ischemic hearts

Because we observed a decrease in *PMIS-C* non-CM proliferation, we asked if there were changes in the status of cardiac fibroblast activation. Following an ischemic injury, CM cell death triggers an immune response and activates the resident cardiac fibroblasts to begin transitioning into a myofibroblast, which have a unique gene signature ^56–59^. To gauge the number of cardiac fibroblasts within the infarct zone, we stained for Vimentin and PDGFRα (Fig. 6A). Results found expression of these pan-fibroblast markers in WT and *PMIS-C* hearts, at 1 and 3 WPI (Fig. 6A). We next analyzed the presence of myofibroblast by examining αSMA expression ^60,61^.

**Fig. 6.**
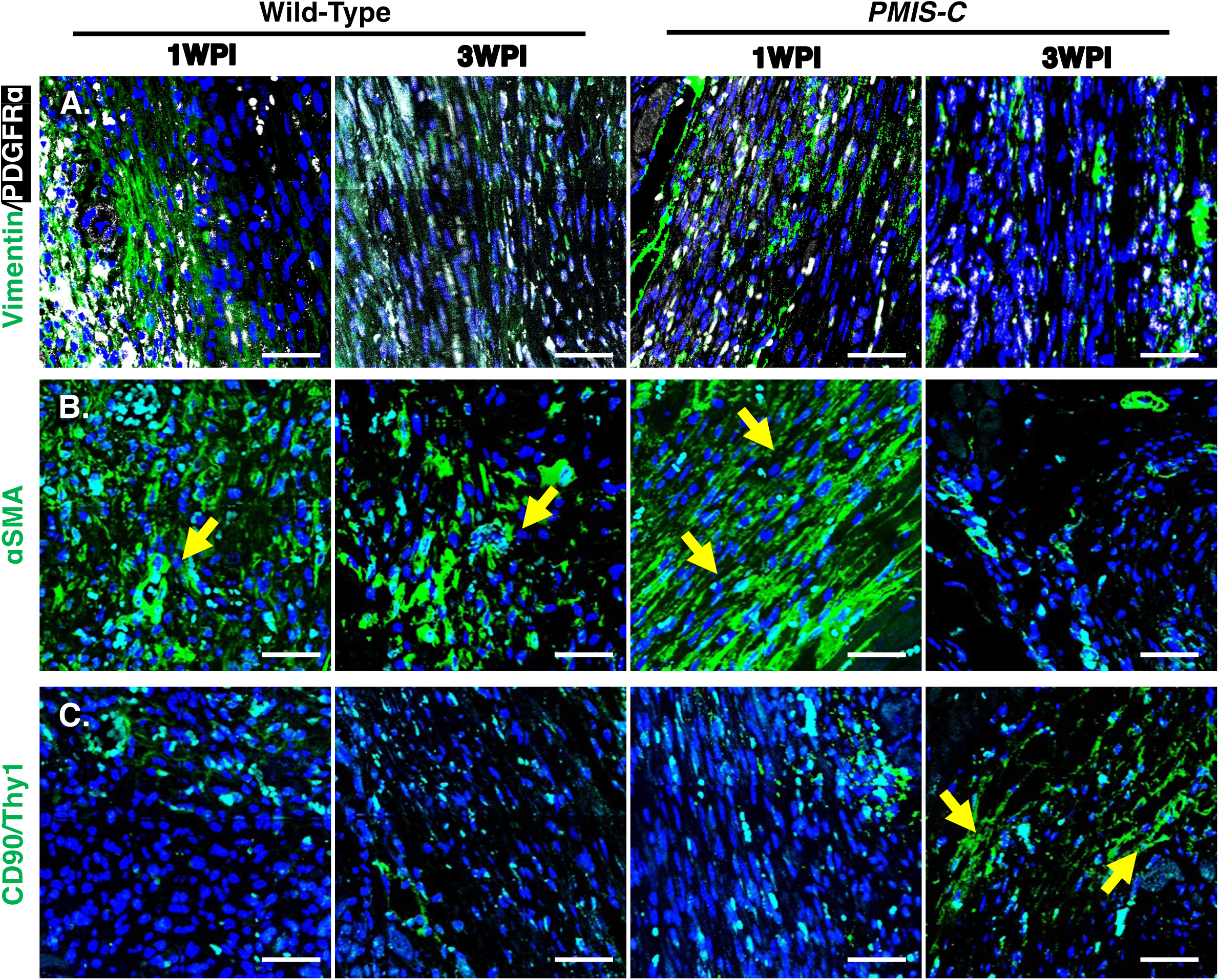
Inhibition of *miR-200c* changes fibroblast gene expression in the ischemic zone. **A)** IF stain for pan-fibroblast markers PDGFRα and Vimentin at 1 and 3 WPI. **B)** Expression of activated myofibroblast cell marker αSMA. **C)** Expression of pro-regenerative cardiac fibroblast marker CD90. Yellow arrows point to cells positive for the respective stain. Scale bar = 50μm.

At 1 WPI, both WT and *PMIS-C* hearts contained αSMA+ cells within the infarct zone (Fig. 6B). However, by 3 WPI the presence of αSMA+ cells were decreased in *PMIS-C* hearts compared to WT. In addition, the presence of CD31+ cells was increased in *PMIS-C* hearts at 3 WPI (Fig. S8). *PMIS-C* hearts also have increased CD90+ cells within the infarct zone at 3 WPI (Fig. 6C). CD90 is thought to mark a subpopulation of fibroblasts that potentially aid in regeneration ^62,63^. We analyzed Collagen I and Fibronectin expression, which are markers of activated fibroblasts, (Fig. S9). Infarct zone expression of these factors was similar at 1 and 3 WPI, in WT and *PMIS-C,* indicating that inhibiting *miR-200c* does not affect the process associated with activated fibroblasts and remodeling of the myocardium following adult injury ^63^.

## DISCUSSION

Our study has found that inhibiting *miR-200c* can promote cardiac repair following an ischemic injury in the adult murine heart. The *PMIS-C* mice showed significantly increased measurements of cardiac function at 3 WPI, which continued to 9 WPI. This functional recovery was accompanied by reduced fibrotic scar size throughout the LV. Mechanistically, *PMIS-C* hearts have increased expression of early heart developmental TFs such as Tbx5, Gata4, and Mef2c, which are *miR-200c* targets ^34^. We also reported an increase in Yap1 in *PMIS-C* murine developing hearts ^34^. Yap has been reported to induce adult CM cell-cycle reentry in adult post-MI hearts ^64^. Our developmental sn-Multiomics analysis identified an immature population of CMs in *PMIS-C* hearts ^34^, and CMs expressing a similar signature are seen following adult cardiac injury. Interestingly, inhibition of *miR-200c* in adult cardiac repair is not limited to CMs, as proliferating non-CMs in the infarct zone were significantly decreased in *PMIS-C*. The rapid heart repair in *PMIS-C* hearts cannot solely be completed by CM “de-differentiation” and proliferation. *PMIS-miR-200c* is ubiquitously expressed, thus inhibiting *miR-200c* across all cell-types in the heart. We hypothesize that decreased *miR-200c* expression in non-CMs is contributing to the repair observed in *PMIS-C* adult mice after a severe MI. Future study will determine the molecular mechanism of non-CMs in ischemic *PMIS-C* hearts.

### miRs in cardiac repair

Studies have shown cardiac repair in model organisms can be achieved through multiple methods and activation of various pathways ^8–15,64–66^. miRs have emerged as an attractive strategy and gene therapy to induce cardiac repair following an ischemic injury of the adult heart. miRs can modulate a variety of transcripts, which can result in cellular changes, leading to repair ^8,19,24^. Overexpression and inhibition of miRs can promote cardiac repair of an ischemic injury. Overexpression of *miR-17-92*, *-199a*, and *-294* can promote CM de-differentiation and proliferation ^8,10,19,67^, leading to regeneration. Inhibition of *miR-34a*, *-99/100*, and *-128* produced comparable results to that of overexpression. ^10,22^. Additionally, expression of a miR cocktail of *miR-1*, *-133*, *-208*, and *-499* in cardiac fibroblasts (CFs) can induce reprogramming into CMs ^13^. Taken together, previous research has suggested that miRs can be an effective approach for promoting cardiac repair.

### The role of *miR-200c* in heart repair

We postulate that the level of adult cardiac repair in *PMIS-C* mice is, in part, due to the unique role and expression profile of *miR-200c*. Previous successes using miRs have focused on ones that maintain a consistent level of expression in adult homeostasis and following cardiac injury. Therefore, these miRs are required to modulate CM cellular functions, and altering their expression could cause unintended effects to the cell.

Second, these miRs are limited in their regulation of transcripts and target pathways, suggesting a conserved and specified role across cell types. In contrast, *miR-200c* has dynamic expression in the embryonic, neonatal, adult heart, and in ischemic injury. We found *miR-200c* was highly expressed in the embryonic heart, while in the adult heart expression is nearly undetectable. As we have shown in other cellular contexts, *miR-200c* expression is required for cell differentiation during development. *miR-200c* inhibits progenitor cell factors to allow for cell differentiation during development ^30,32,34,38,68^. Following MI, expression of *miR-200c* increases, suggesting a role in modulating the disease state. In terms of targets, *miR-200c* is known to regulate stem cell factors (Sox2, Klf4) ^32,69^, Hippo ^70^, TGFβ/BMP/Smad ^38^, FGF ^71^, Wnt/β-Catenin ^72^, EMT ^73^, and hypoxia/Vegf signaling ^74^, all of which have been implicated in assisting cardiac regeneration across various cell types ^4^.

Moreover, in cardiac development, we found that *miR-200c* directly targets *Tbx5*, *Gata4*, and *Mef2c* transcripts. These factors in CMs, post-MI, are thought to promote de-differentiation ^12,15,22^. Taken together, *miR-200c* has a broad range of pathway targets and specific expression profiles in the heart, aiding the enhanced rate of adult repair seen in *PMIS-C* animals. In support of our data, another study showed a similar response by the removal of the Hippo pathway component *Salvador*, which, recovered cardiac function by 3-4 weeks ^75^.

### Contribution of existing CMs to the repaired myocardium

The results from the proliferation assay suggest that repair of the new myocardium, in part, arises from the proliferation of pre-existing CMs of the BZ ^46^. We identified immature CMs in the developing heart of *PMIS-C* mice, and these cells could contribute to part of the new adult myocardium in the injured *PMIS-C* hearts ^34^. However, this proliferation is likely not sufficient to produce the whole of the myocardium within the repaired *PMIS-C* hearts. We also found a population of CD90+ cells in the *PMIS-C* infarct zone at 3 WPI, not observed in WT mice. These CD90+ cells mark mesenchymal stem cells and cardiac fibroblasts that may be beneficial for heart repair ^76^. Future studies will look to define the populations of cells that produce the new myocardium in *PMIS-C* hearts (Fig. 7).

**Fig. 7.**
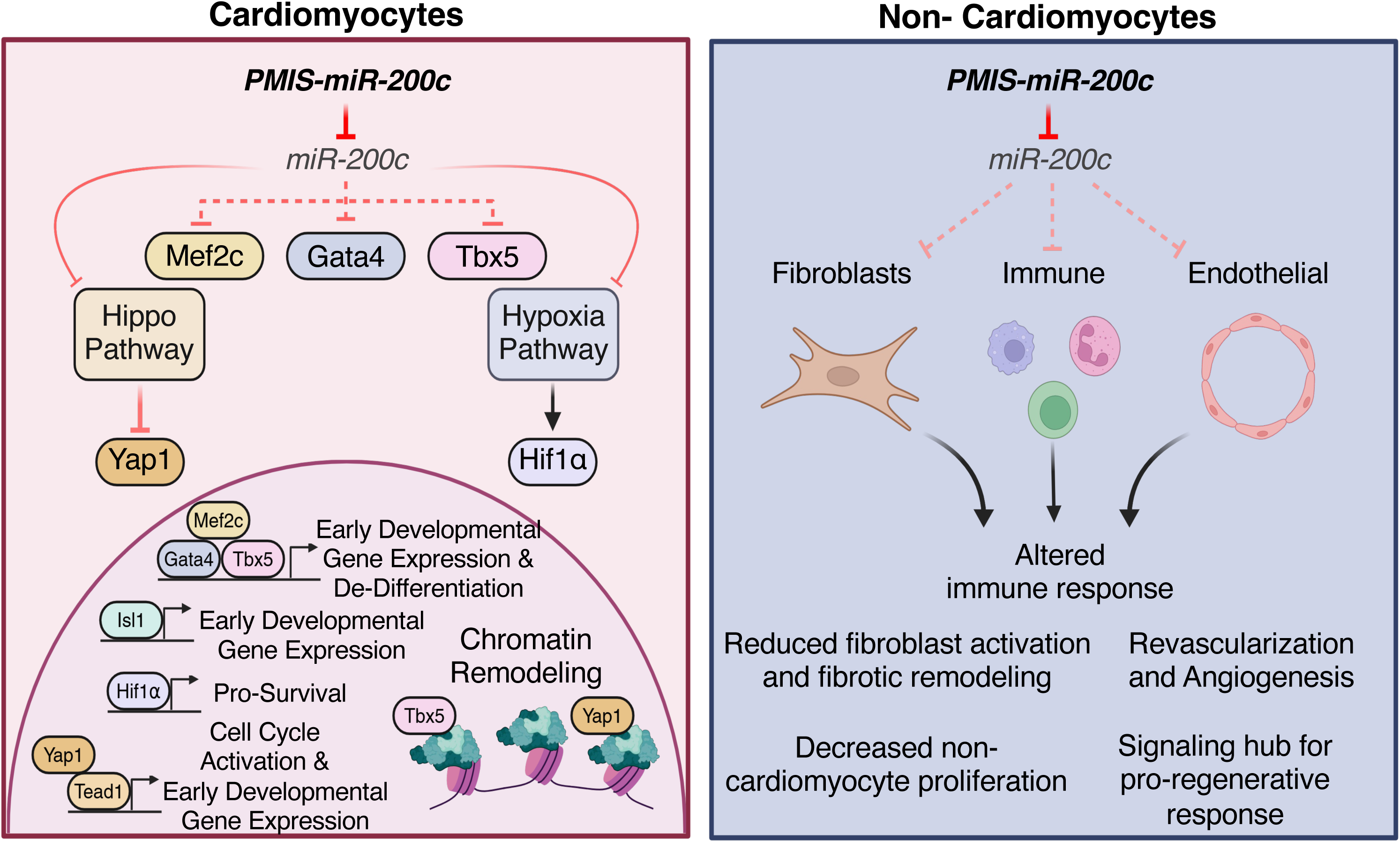
*PMIS-miR-200c* mechanisms for heart regeneration and repair. Different cell types within the heart may be affected by the inhibition of *miR-200c*.

### Contribution of non-CMs to the repaired myocardium

It is well established that non-CM cell-types have an essential role following an ischemic injury ^63,77–80^. *miR-200c* is known to be expressed in non-CMs of the heart ^34^, suggesting a role for this miR in multiple lineages. As mentioned, *miR-200c* can target factors across a range of pathways, including hypoxia (Fig. 7), which is activated following injury. Future experiments will look to understand which cell types are affected and the molecular targets of *miR-200c* in these cells.

### Limitations of this study

snRNA multiomics of *PMIS-miR-200c* (*PMIS-C*) mice revealed an immature cardiomyocyte cell state ^34^, and these immature CMs may offer a protective role against ischemic injury. However, all mice undergo apoptosis after an MI and form fibrotic lesions, with the presence of activated fibroblasts. Cell proliferation assays show that EdU retaining cells are low and similar between WT and *PMIS-C* mice after 3 weeks indicating both mice had rapid CM proliferation in the border zone. However, the increased levels of BrdU labeling in the *PMIS-C* mice after 1 week compared to WT indicate continued CM proliferation in *PMIS-C* mice. We speculate the increase in CM’s in the *PMIS-C* infarct zone reduced fibrosis, while the WT mice had an increase in non-cardiomyocyte proliferation. Further studies will require direct injection of *PMIS-miR-200c* to WT mice after an MI, to observe potential therapeutic effects and differentiate protective effects. The results reported here provide direct evidence that *miR-200c* inhibition may repair the heart after a severe ischemic injury.

## ACKNOWLEDGMENTS

We thank members of the Amendt laboratory for their expertise and helpful discussions and previous lab members for contributing to the study. We acknowledge contributions from Kathy Zimmerman and Ariana Batz for echocardiograms and analyses, William Kutschke for mouse LAD surgeries, the University of Iowa Cardiovascular Research Center and Cardiovascular Phenotyping Core for advice and support of this project. We thank the University of Iowa Cardiovascular research group for great discussions and comments on this study. We also thank the University of Iowa, Center for Craniofacial Anomalies for their support.

## Sources of Funding

This work was supported by funding from, National Institutes of Health grant R01DE028527 and R43DE027569, and the University of Iowa to BAA; National Institutes of Health grant S10OD038119 to the University of Iowa, Cardiovascular Phenotyping Core.

## Author Contributions

R.L., performed the experimental studies, analyzed data and prepared figures and a draft of the manuscript; M.S., performed experimental studies, prepared figures, and edited the manuscript; S.E., performed the experimental studies, analyzed data and managed funds and reagents; W.K., performed mouse LAD experiments, analyzed data, edited the manuscript, provided critical input on experimental design; B.A.A., project development, funding, analyzed data, prepared the manuscript.

## Competing Interests

Brad A. Amendt is the CSO and founder of NaturemiRI, LLC.

All other authors have no disclosures to report.

## Data and materials availability

All data are available in the main text or the supplementary materials.

## SUPPLEMENTAL FIGURE LEGENDS

**Fig. S1. MI size and ejection fraction in heart failure. A)** Ejection fraction of all mice that met the criteria of inclusion in the MI study 1 DPI (see Materials and Methods). **B)** Infarct zone fraction of the same group of mice. The statistical test performed was a two-tailed t-test. **C)** Kaplan-Meier death curve of WT, *PMIS-A*, and *PMIS-C* mice subject to MI. *PMIS-C* mice survival rate was significantly improved compared to WT and *PMIS-A*. No mice died after 7 DPI. The statistical test performed was a Log-rank test.

**Fig. S2. *PMIS-C* mice show functional recovery post-MI and normal heart rates. A)** Delta IZ (infarct zone) of mice that received MI and survived to 3 WPI and IZ fraction of WT, *PMIS-A*, and *PMIS-C* at 3 WPI. **B)** End-diastolic volume and end-systolic volume of WT, *PMIS-A*, and *PMIS-C* mice subject to MI or sham procedure at 3 WPI. **C)** Heart rate of mice subject to MI from 1 DPI to 9 WPI. The statistic test performed was one-way ANOVA with multiple comparisons. The statistical test performed was a two-tailed t-test.

**Fig. S3. MI results in cell death in the infarct zone across all genotypes. A)** IF stain of left ventricle section in the infarct zone of WT, *PMIS-A,* and *PMIS-C* for Cleaved Caspase 3 (CC3) (Red), cTnT (White), and Vimentin (Vim, Green) at 3 DPI. Expression of CC3 is seen in all the hearts, indicating cell death following MI. Scale bar = 100μm. **B)** Trichrome staining of transverse ventricle sections of WT, *PMIS-A*, and *PMIS-C* mice subject to MI, 1 WPI. **C-D)** Magnified image of the infarct zone (IZ) **(C)** and border zone (BZ) **(D)**. Blue stain is seen in all three hearts, indicating scar formation. Scale bar = 500 μm **(B)**. Scale bar = 100 μm **(C-D)**.

**Fig. S4. *PMIS-C* hearts have minimal fibrotic scarring at 9 WPI. A)** Trichrome staining of transverse ventricle sections of individual WT, *PMIS-A*, and *PMIS-C* subject to MI, 9 WPI. Echocardiogram LVEF for each heart is indicated above. Scale bar = 1mm.

**Fig. S5. Expression of cardiogenic TFs post-MI. A-B)** Expression of *Tbx5*, *Gata4*, and *Nkx2.5* at 1WPI (A) and 3 WPI (B) in WT, *PMIS-A*, and *PMIS-C.* RNA was isolated from whole ventricle tissue. The statistical test performed was a two-tailed t-test *=p≤0.05, **=p≤0.01, ***=p≤0.001.

**Fig. S6. Nppa and Sox5 expression in CMs of injured hearts. A-B)** IF stain of border zone CMs for Nppa (A) and Sox5 (B), in WT and *PMIS-C* sham/uninjured or at 3 WPI. Quantitation of Nppa/cTnT+ and Sox5+ cells are shown. Scale bar = 50μm. The statistical test performed was a two-tailed t-test.

**Fig. S7. Nppa+/Sox5+ cells in the border zone and single-positive cells are found in the regeneration zone post-MI. A-B)** IF stain of left ventricle sections for Nppa (Red), Sox5 (Green), and cTnT (White) in WT (A) and *PMIS-C* (B) in the BZ, IZ, and RZ at 1 WPI. Left: Nppa+/Sox5+ CMs are found at the border zone in *PMIS-C* hearts (Arrow). Center: Nppa+ CMs are found in the infarct zone, and Sox5 expression appears in cells most likely of epicardial origin. Right: Nppa+ and Sox5+ cells are seen in the remote zone in proximity to vasculature structures (Arrow). The presence of these cells increased in *PMIS-C* hearts. Scale bar = 50μm.

**Fig. S8. *PMIS-C* heart has vascular structures within the RegZ. A-B**) IF stain of left ventricle sections CD31 (Green) and cTnT (Red) in WT and *PMIS-C* in sham or at 3 WPI. Representative border zone images show vasculature present in sham hearts. *PMIS-C* heart at 3 WPI, CD31+ endothelial cells. Scale bar = 100μm.

**Fig. S9. Expression of activated fibroblast markers in injured hearts A-B)** IF stain for activated fibroblast markers Collagen I and Fibronectin in WT and PMIS-C hearts at 1 and 3 WPI. Scale bar = 50μm.

## Notes

### Competing Interest Statement

Brad Amendt is CSO of NaturemiRI

