## Supplementary material for "Activation of developmental transcription factors using RNA technology promotes heart repair": Suppl Files

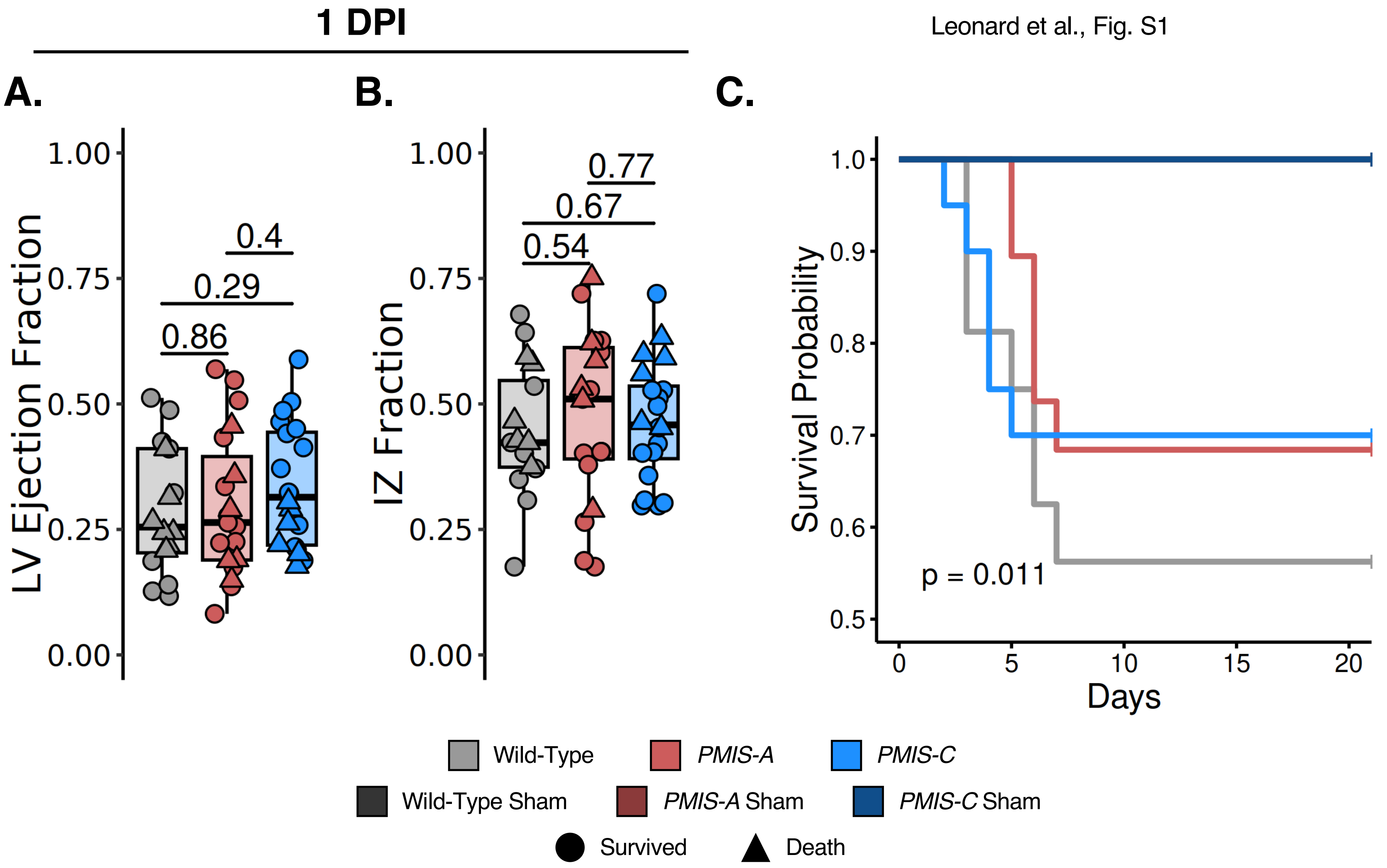


**Fig. S1. MI size and ejection fraction in heart failure. A)** Ejection fraction of all mice that met the criteria of inclusion in the MI study 1 DPI (see Materials and Methods). **B)** Infarct zone fraction of the same group of mice. The statistical test performed was a two-tailed t-test. **C)** Kaplan-Meier death curve of WT, *PMIS-A*, and *PMIS-C* mice subject to MI. *PMIS-C* mice survival rate was significantly improved compared to WT and *PMIS-A*. No mice died after 7 DPI. The statistical test performed was a Log-rank test.

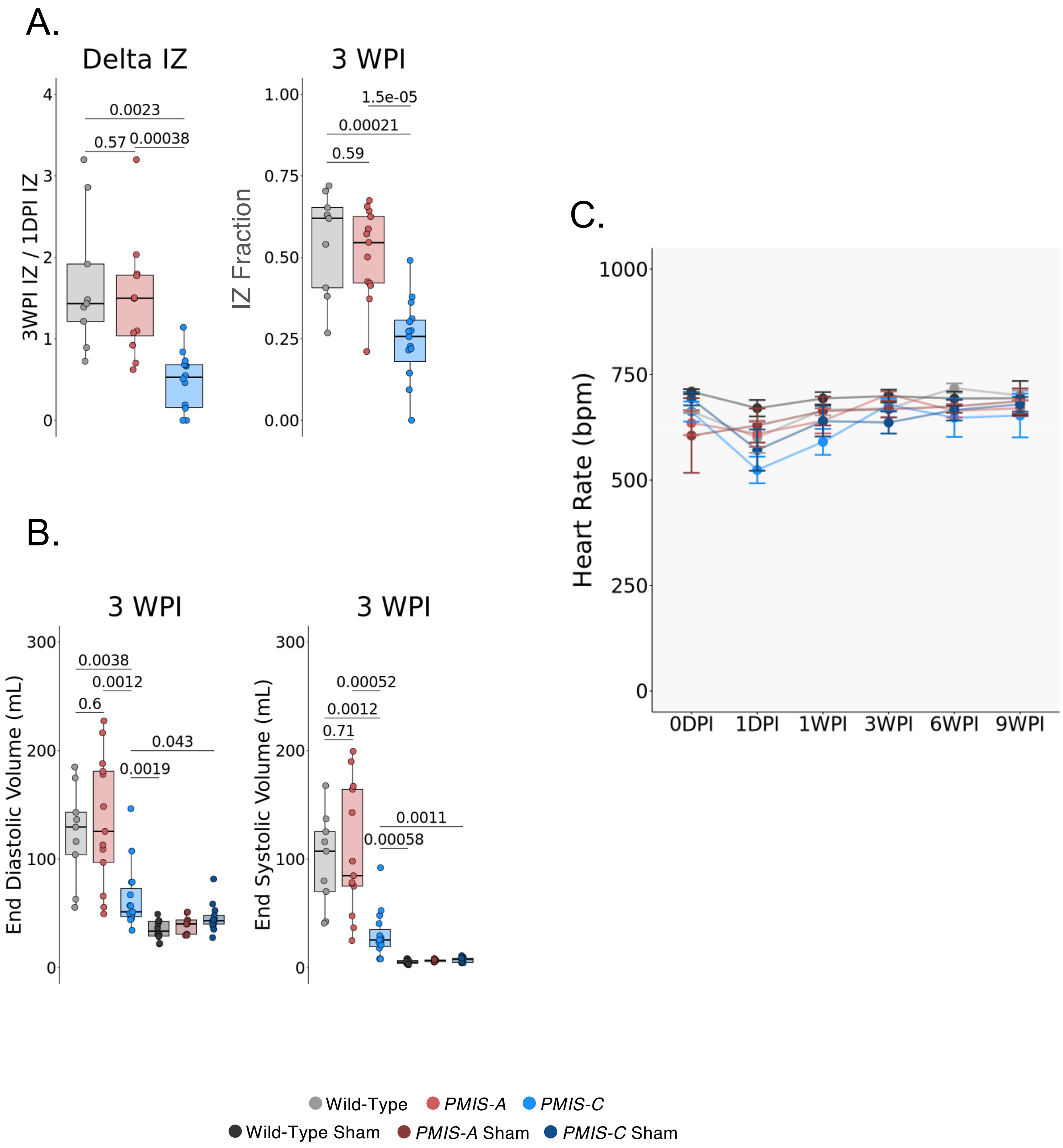


**Fig. S2. *PMIS-C* mice show functional recovery post-MI and normal heart rates. A)** Delta IZ (infarct zone) of mice that received MI and survived to 3 WPI and IZ fraction of WT, *PMIS-A*, and *PMIS-C* at 3 WPI. **B)** End-diastolic volume and end-systolic volume of WT, *PMIS-A*, and *PMIS-C* mice subject to MI or sham procedure at 3 WPI. **C)** Heart rate of mice subject to MI from 1 DPI to 9 WPI. The statistic test performed was one-way ANOVA with multiple comparisons. The statistical test performed was a two-tailed t-test.

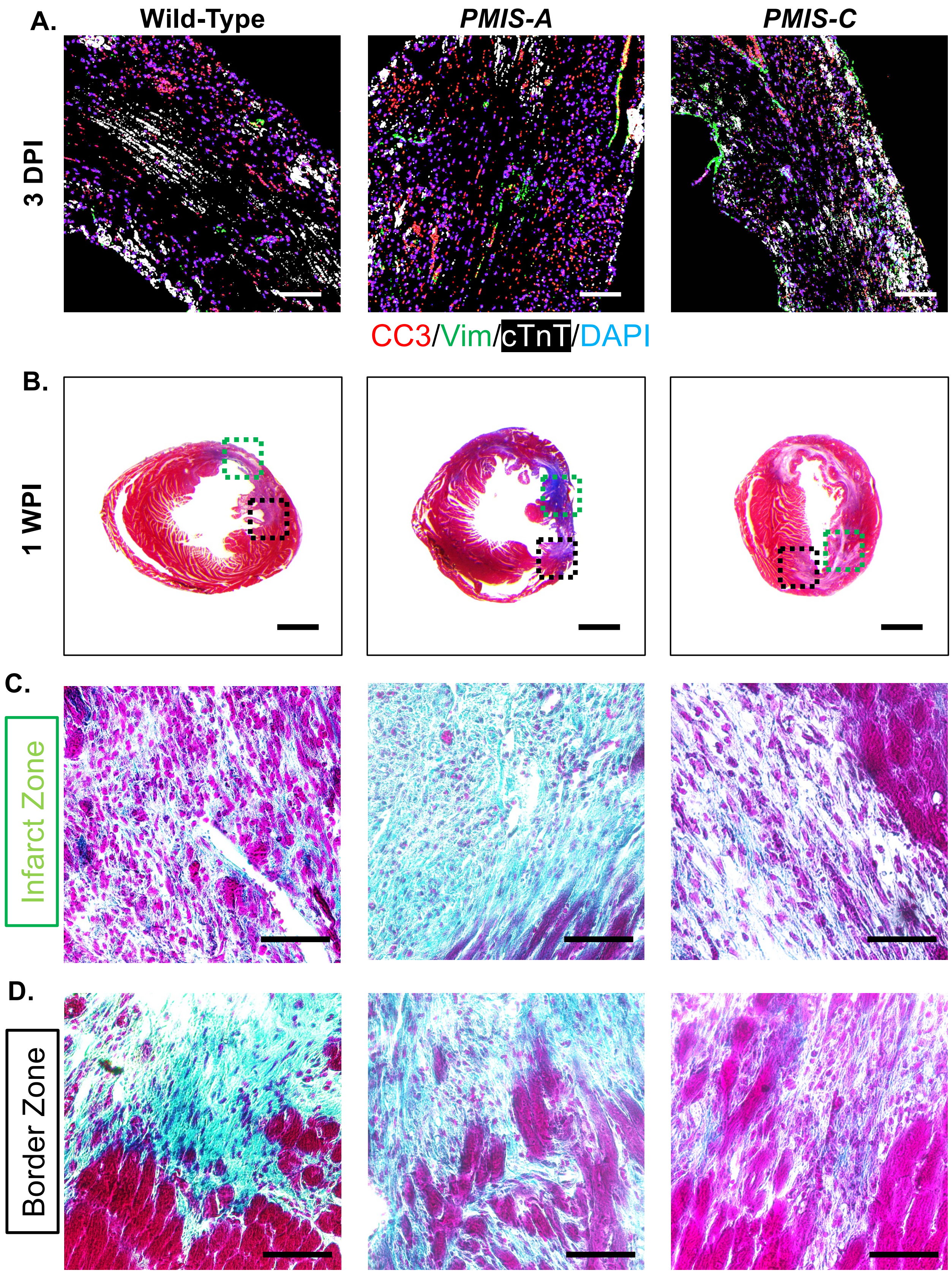


**Fig. S3. MI results in cell death in the infarct zone across all genotypes. A)** IF stain of left ventricle section in the infarct zone of WT, *PMIS-A,* and *PMIS-C* for Cleaved Caspase 3 (CC3) (Red), cTnT (White), and Vimentin (Vim, Green) at 3 DPI. Expression of CC3 is seen in all the hearts, indicating cell death following MI. Scale bar = 100µm. **B)** Trichrome staining of transverse ventricle sections of WT, *PMIS-A*, and *PMIS-C* mice subject to MI, 1 WPI. **C-D)** Magnified image of the infarct zone (IZ) **(C)** and border zone (BZ) **(D)**. Blue stain is seen in all three hearts, indicating scar formation. Scale bar = 500 µm **(B)**. Scale bar = 100 µm **(C-D)**.

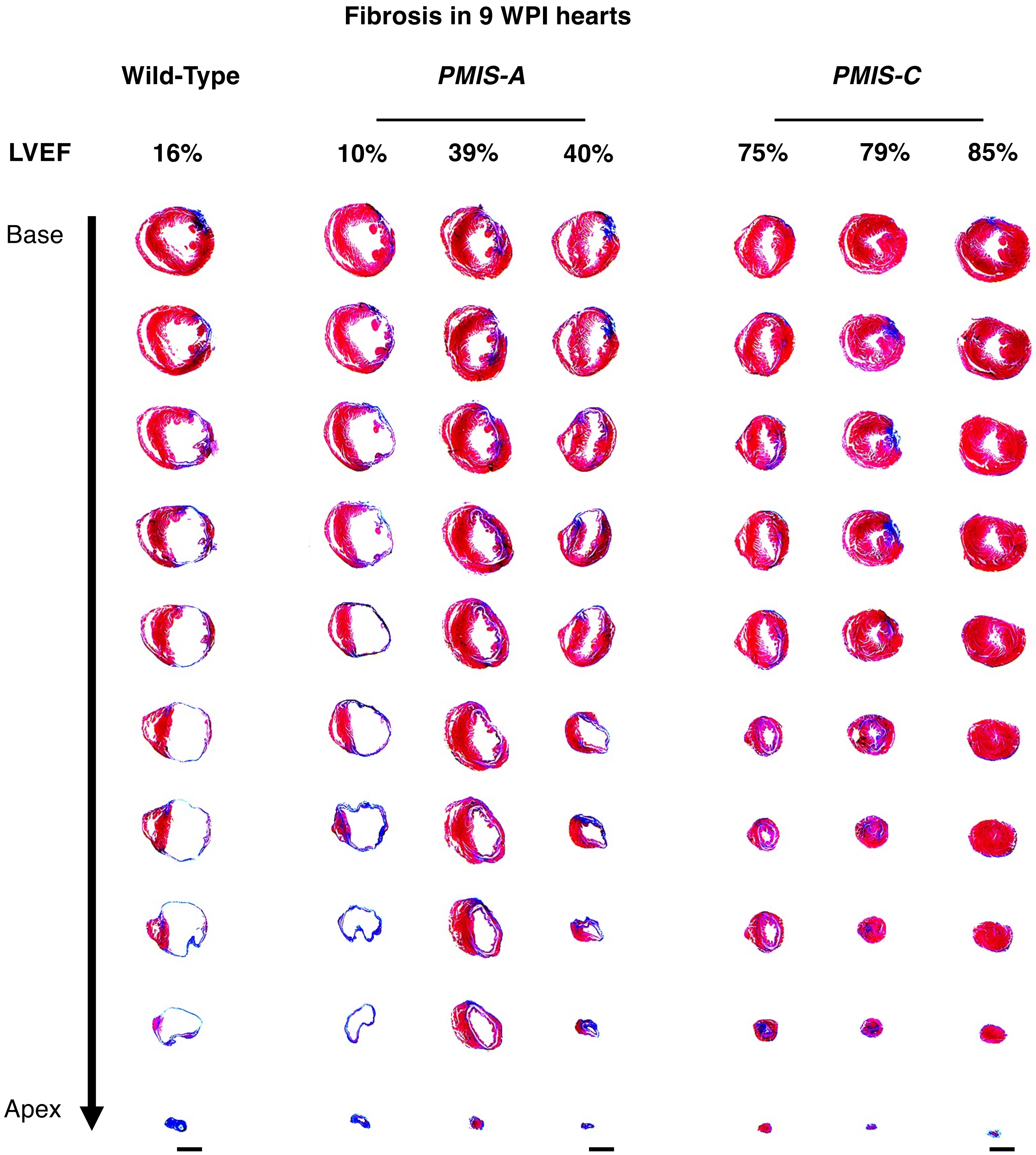


**Fig. S4.** ***PMIS-C*** **hearts have minimal fibrotic scarring at 9 WPI. A)** Trichrome staining of transverse ventricle sections of individual WT, *PMIS-A*, and *PMIS-C* subject to MI, 9 WPI. Echocardiogram LVEF for each heart is indicated above. Scale bar = 1mm.

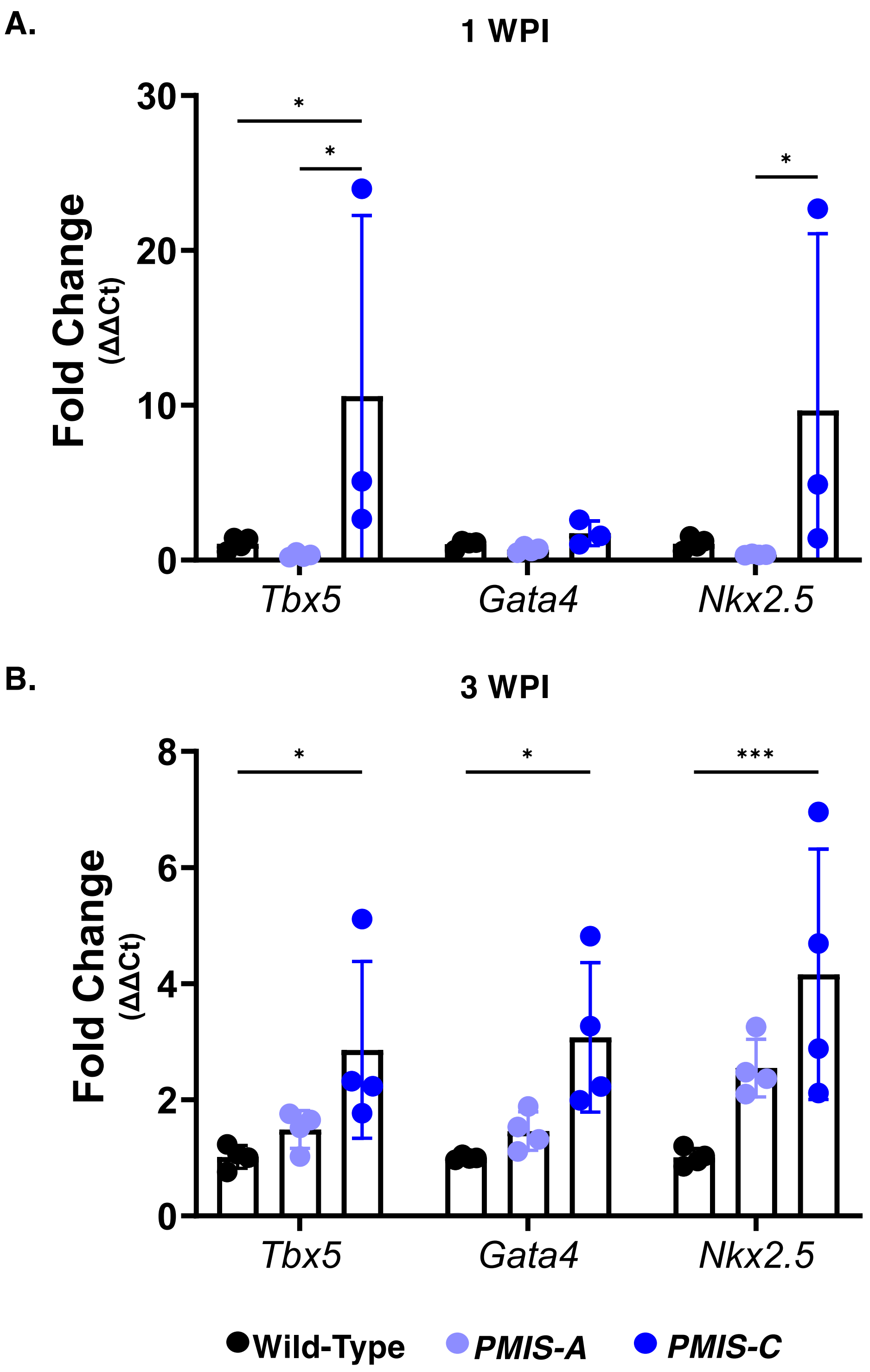


**Fig. S5. Expression of cardiogenic TFs post-MI. A-B)** Expression of *Tbx5*, *Gata4*, and *Nkx2.5* at 1WPI (A) and 3 WPI (B) in WT, *PMIS-A*, and *PMIS-C*. RNA was isolated from whole ventricle tissue. The statistical test performed was a two-tailed t-test $*=p\leq0.05$, $**=p\leq0.01$, $***=p\leq0.001$.

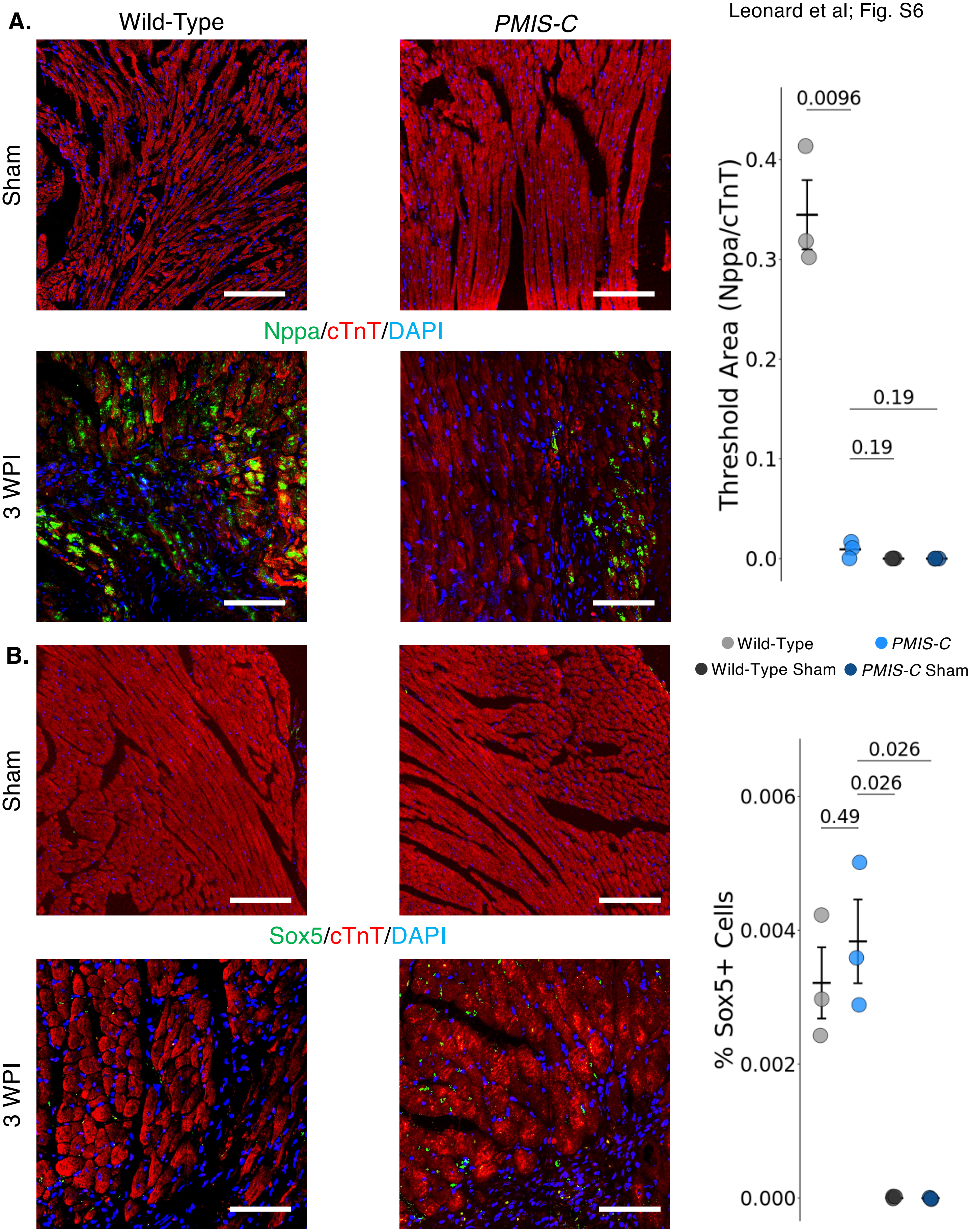


**Fig. S6. Nppa and Sox5 expression in CMs of injured hearts. A-B)** IF stain of border zone CMs for Nppa (A) and Sox5 (B), in WT and *PMIS-C* sham/uninjured or at 3 WPI. Quantitation of Nppa/cTnT+ and Sox5+ cells are shown. Scale bar = 50µm. The statistical test performed was a two-tailed t-test.

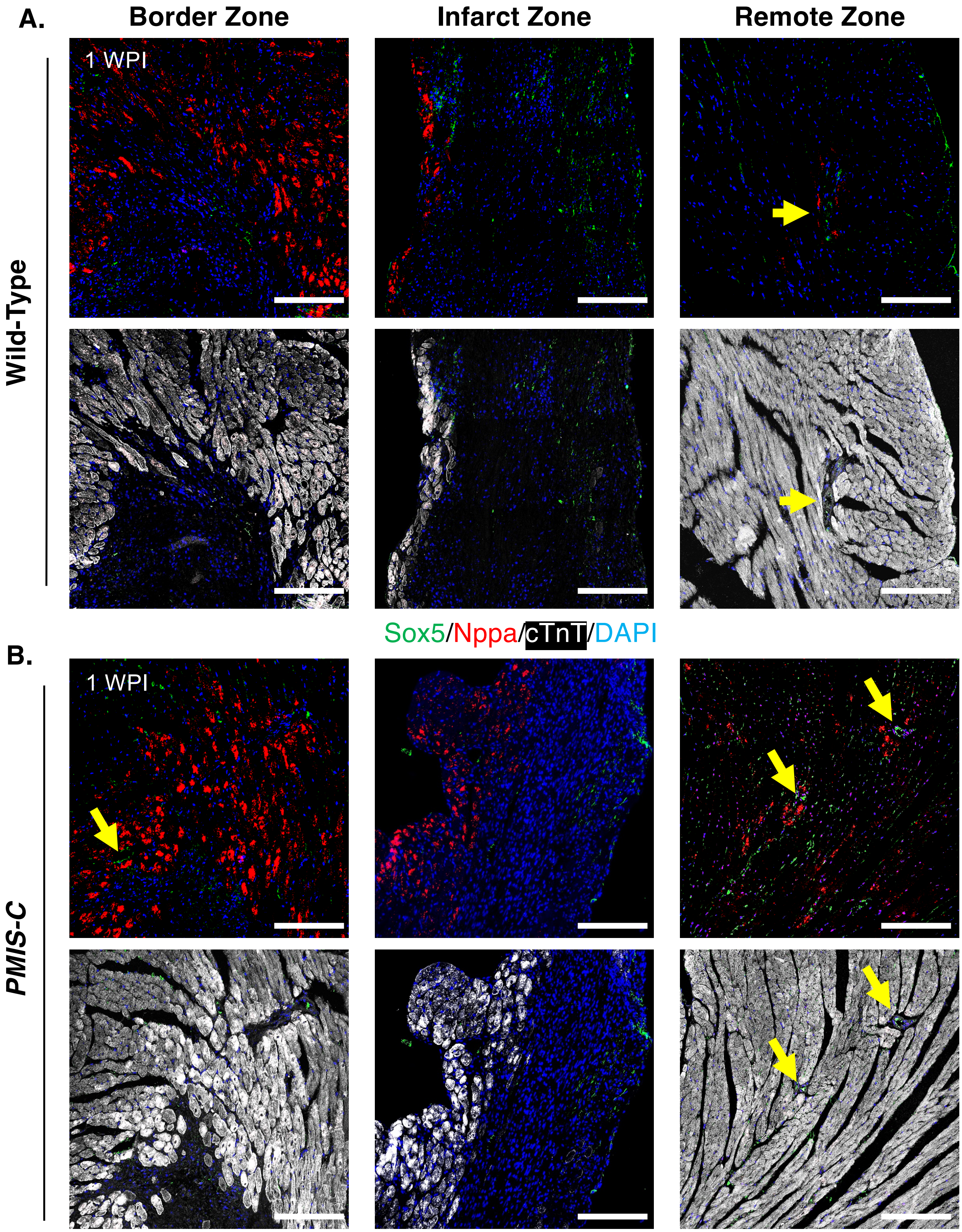


**Fig. S7. Nppa+/Sox5+ cells in the border zone and single-positive cells are found in the regeneration zone post-MI. A-B)** IF stain of left ventricle sections for Nppa (Red), Sox5 (Green), and cTnT (White) in WT (A) and *PMIS-C* (B) in the BZ, IZ, and RZ at 1 WPI. Left: Nppa+/Sox5+ CMs are found at the border zone in *PMIS-C* hearts (Arrow). Center: Nppa+ CMs are found in the infarct zone, and Sox5 expression appears in cells most likely of epicardial origin. Right: Nppa+ and Sox5+ cells are seen in the remote zone in proximity to vasculature structures (Arrow). The presence of these cells increased in *PMIS-C* hearts. Scale bar = 50µm.

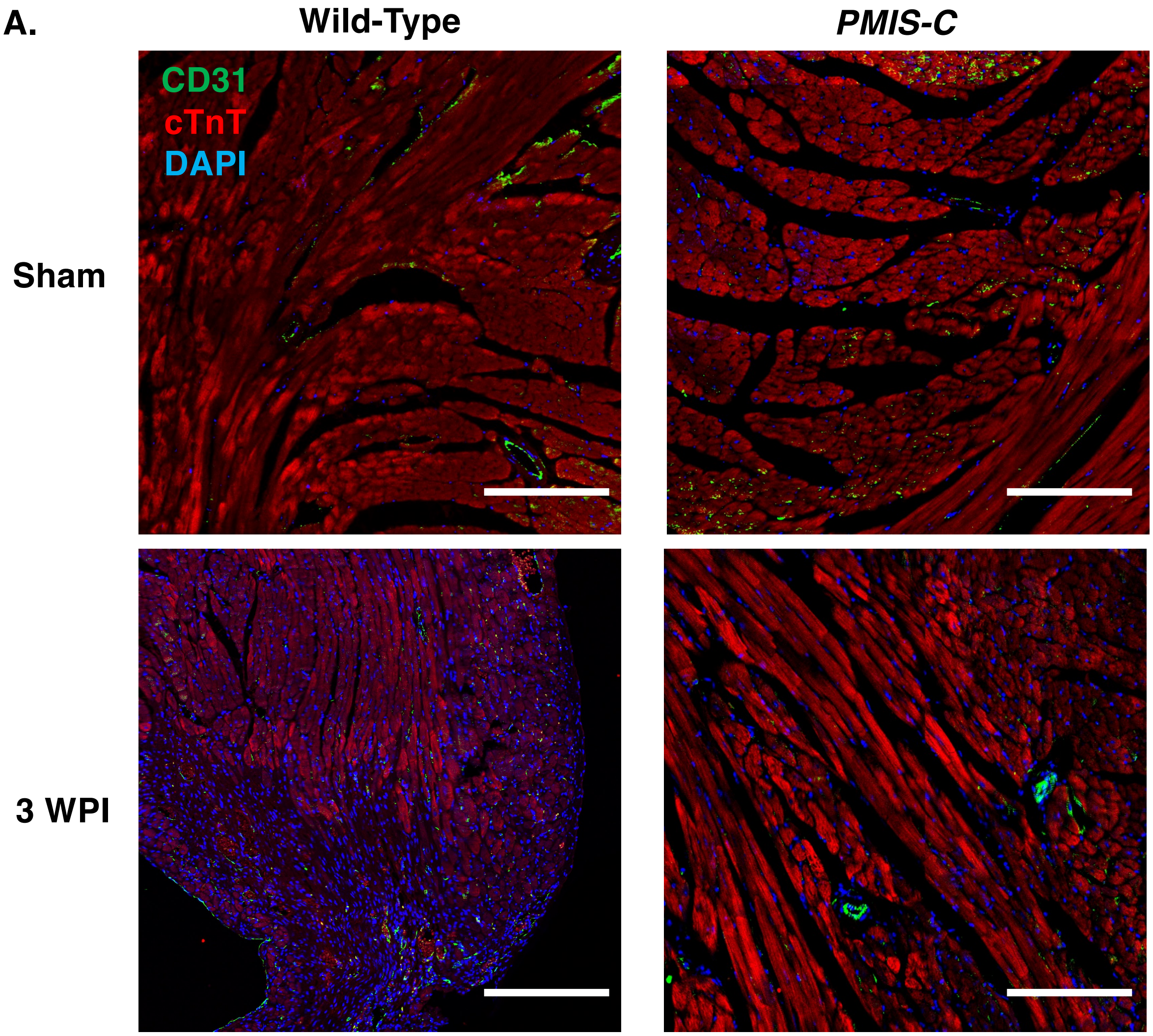


**Fig. S8. *PMIS-C* heart has vascular structures within the RegZ. A-B**) IF stain of left ventricle sections CD31 (Green) and cTnT (Red) in WT and *PMIS-C* in sham or at 3 WPI. Representative border zone images show vasculature present in sham hearts. *PMIS-C* heart at 3 WPI, CD31+ endothelial cells. Scale bar = 100µm.

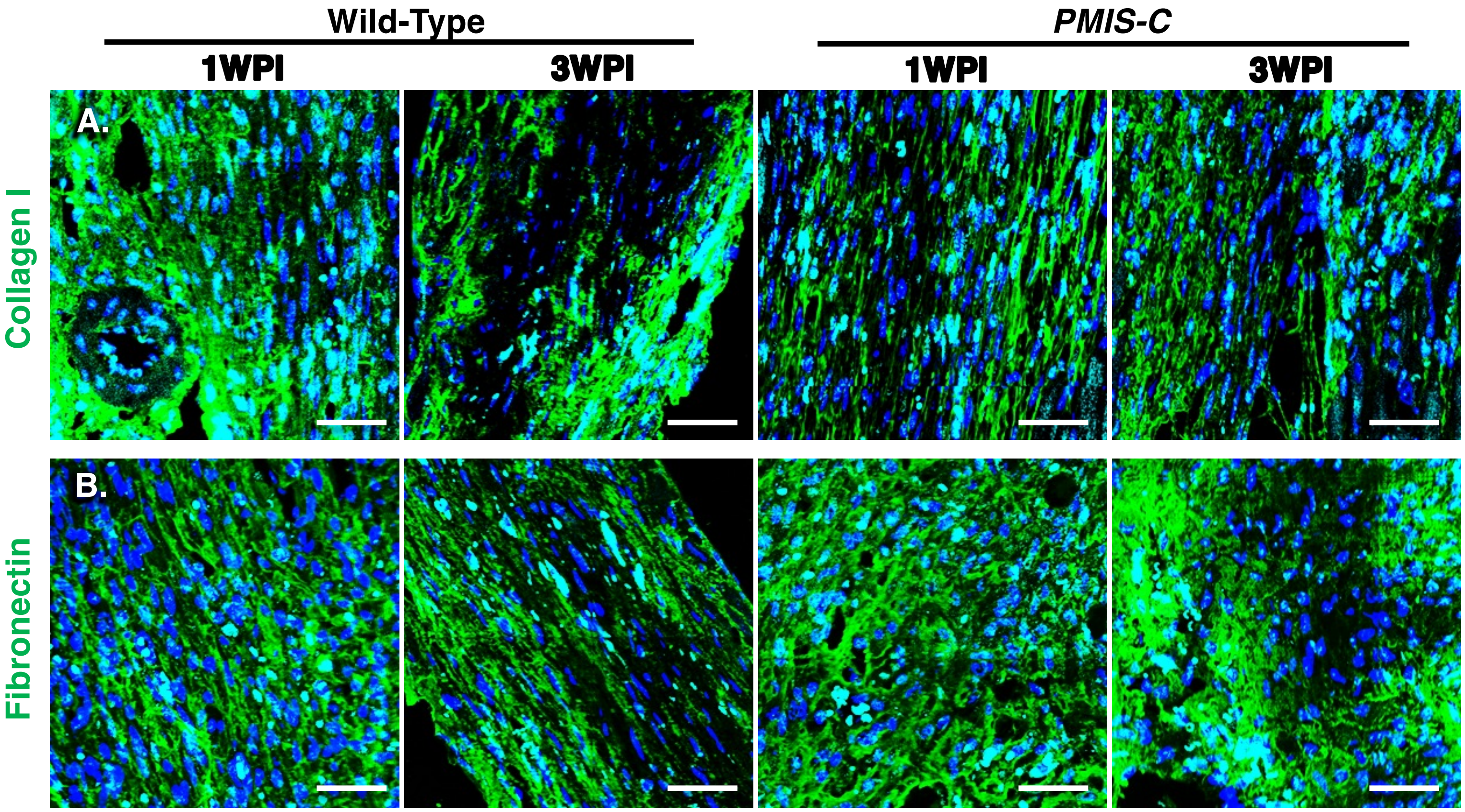


**Fig. S9. Expression of activated fibroblast markers in injured hearts A-B)** IF stain for activated fibroblast markers Collagen I and Fibronectin in WT and PMIS-C hearts at 1 and 3 WPI. Scale bar = 50μm.
